# Multi-omics features-based machine learning method improve immunotherapy response in clear cell renal cell carcinoma

**DOI:** 10.1101/2023.11.24.568360

**Authors:** Yiqun Zhang, Zhihua Pei

**Affiliations:** independent researcher; Hubei Key Laboratory of Agricultural Bioinformatics, College of Informatics, Huazhong Agricultural University

## Abstract

Programmed cell death 1 (PD-1) or PD-ligand 1 (PD-L1) blocker-based strategies have improved the survival outcomes of clear cell renal cell carcinomas (ccRCCs) in recent years, but only a small number of patients have benefited from them. In this study, we identified three inflammatory features through over 1900 autoimmune nephropathy patients-related bulk RNA sequencing, single-cell RNA sequencing analysis, and three immunogenic signatures using genomics (TIs), both of which are associated with response to immune checkpoint blocks (ICBs) and the survival of ccRCC patients. Here, we developed a framework with a TIs-based machine learning approach to accurately predict ICB efficacy. We enrolled more than 1000 ccRCC patients with ICB treatment from five cohorts to apply the model and demonstrated its excellent specificity and robustness. Moreover, our model outperforms well-known ICB predictive biomarkers such as tumor mutational burden (TMB), PD-L1 expression, and tumor immune microenvironment (TME). Overall, the TIs-ML model provides a novel method for guiding precise immunotherapy in ccRCC.

## Introduction

Immune checkpoint blockades (ICBs) that block programmed death 1 (PD-1), programmed cell death ligand 1 (PD-L1), or cytotoxic T lymphocyte antigen 4 (CTLA4) have ushered in a new era in the treatment of advanced cancer ^1,2^. Notably, PD-(L)1-based therapies have become the standard-of-care option for advanced clear cell renal cell carcinomas (ccRCCs), regardless of the number of prior lines of therapy received^3-7^. However, only a subset of ccRCC patients (20%–50%) respond to immune monotherapy or combination therapies^3,4,8,9^. Therefore, a comprehensive approach for identifying biomarkers that are associated with potential immunotherapy responders or survival-benefit patients is a significant unmet medical need in ccRCC with ICB treatment.

The major challenge is to find comprehensive biomarkers that can predict response and survival to immunotherapy alone or in combination for ccRCC, regardless of lines of therapy. For example, PD-L1 expression, which belongs to proteomics, and tumor mutation burden (TMB), which is defined by genomic alterations, both are predictive biomarkers of response to PD-1/PD-L1 inhibitors in non-small cell lung cancer, melanoma, colorectal cancer, and head and neck cancers^10-16^, but applying them to predict the efficacy of immunotherapy for ccRCC undergoing ICB therapy has been continuously questioned^17,18^. Bulk RNA sequencing (RNA-seq) or single-cell RNA sequencing (scRNA-seq)-based multi-gene prognostic models and comprehensive somatic alteration signatures through genomics analysis are based on one single omics, improving the accuracy of ICB response prediction to some extent^17-20^. Indeed, recent studies have demonstrated that immune infiltration interacts with genomic features in ccRCC^21,22^. Multi-omics-based machine learning (ML) models outperform single-omics methods in predicting therapy efficacy, which has been proven in esophageal cancer and breast cancer^23,24^. These biomarkers identified in patients with ICB can only partially explain the response to immunotherapy treatment^25^, indicating that a more integrated multi-omics biomarker is needed for ccRCC patients.

Nonetheless, the novel multi-omics features need to be investigated not only from immunogenic characteristics in oncological patients with ICB, but also from inflammatory profiles in non-oncological patients with autoimmune disease. The kidney has a more complex immune environment than other organs, which is vulnerable to autoimmune attacks^26,27^. There is increasing evidence that the inflammatory signatures of autoimmune nephropathy are strongly correlated with the efficacy of immunotherapy for patients with advanced malignant tumors, especially in ccRCC^28,29^. The potential reason is that immune-related adverse effects (irAEs) derived from immunotherapy, which can predict the prognosis of immunotherapy, are similar in pathogenesis and clinical therapy to autoimmune-related diseasesy^30-33^. However, there is a lack of relevant literature on integrating ML-based inflammatory and immunogenic multi-omics approaches to improve the predictive specificity and robustness of ccRCC with ICB regimens.

In this present research, we offer an inflammatory and immunogenic-based (TIs) multi-omics ML framework that can (i) generate robust predictions about response and survival on ICB datasets, (ii) construct predictions specifically for ccRCC with ICB, and (iii) develop overall survival predictions in ccRCC. Specifically, we identified three types of inflammatory biomarkers in more than 1900 patients with autoimmune nephropathy using RNA-seq and scRNA-seq data, and three types of immunogenic signatures by genomics analysis in ccRCC with immunotherapy. Ultimately, with the TIs-ML model, we can reliably distinguish responders or survival-benefit patients in over 1000 ccRCCs with ICB. Predictive models based on individual 6-type features have presented a great variation in performance, while the ensemble model outperforms either individual classifier. The performance of the aggregate model was superior to the biomarkers identified in immunotherapy-treated patients, such as tumor mutational burden (TMB), PD-L1 expression, and tumor immune microenvironment (TIME). Moreover, we obtained 716 genes related to inflammation in autoimmune nephropathy, which were enriched in the largest pathway as “regulation of lymphocyte activation”. Using these genes, we clustered ccRCC into two categories, which have different treatment responses and prognosis preferences. As a result, we find that the inflammatory profiles of immune-mediated kidney diseases are strongly associated with the outcome of immunotherapy in ccRCC, and our integrated model improves the prediction of the ICB response and survival.

## Result

### Overview of TIs-ML method for ICB predictions

Previously published works suggested a connection between the effectiveness of immunotherapy and autoimmune diseases^28,29^. The workflow of this study is shown in Fig. 1. We adopted five autoimmune nephropathies^34,35^ with scRNA or RNA-seq data. Ultimately, we investigated a total of 42 cohorts, including eight lupus nephritis (LN) cohorts, nine IgA nephropathy (IgAN) cohorts, twelve membranous nephropathy (MN) cohorts, nine focal segmental glomerulosclerosis (FSGS) cohorts, and four antineutrophil cytoplasmic antibody vasculitis (ANCA-AAV) cohorts, which were over 1900 samples. We obtained a total of 5 cohorts of ccRCC with ICB treatment, which included over 1000 patients.

**Fig. 1.**
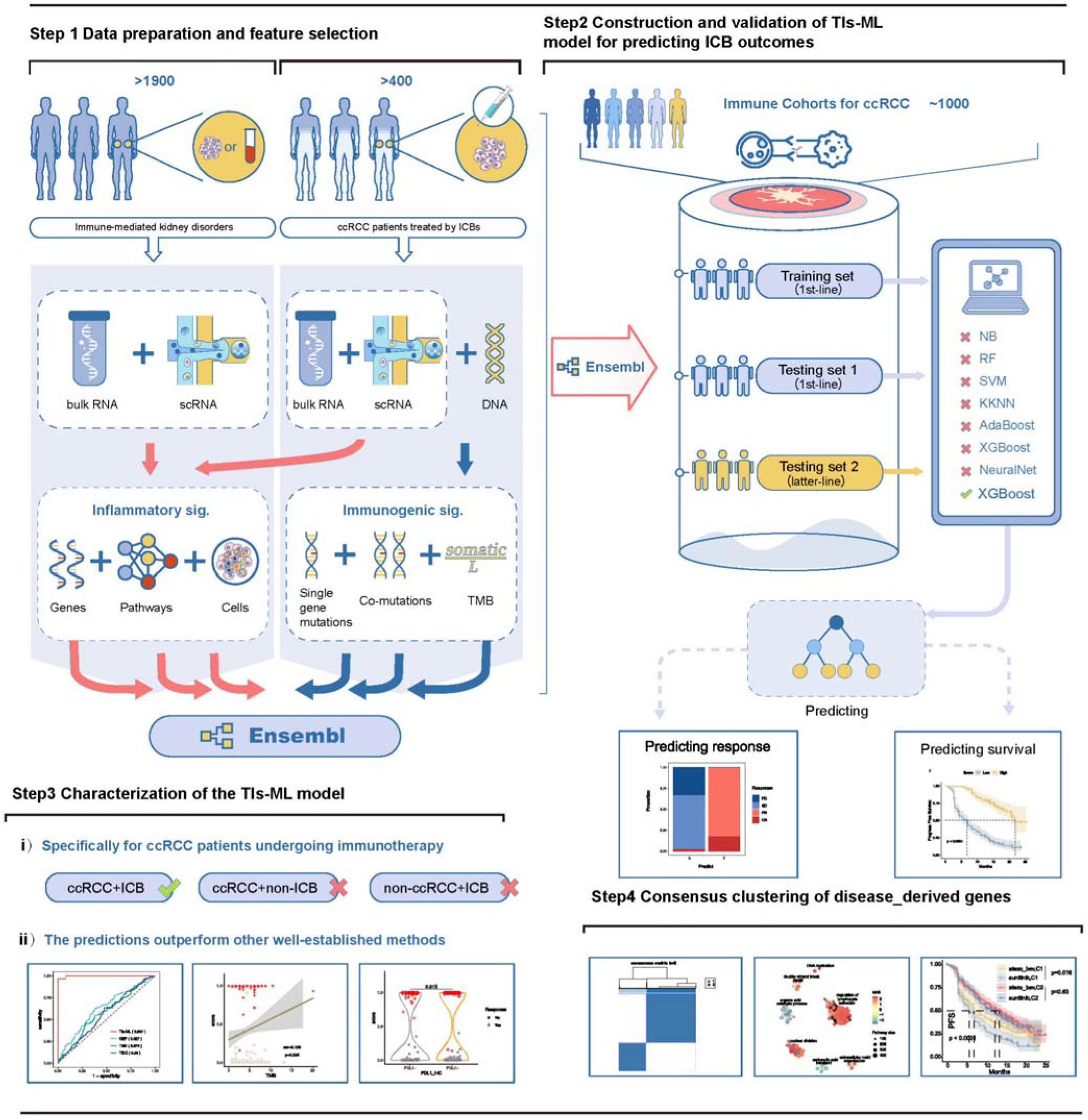
Workflow of this work. Tissue or plasma samples were collected from multi-studies that enrolled more than 1900 patients diagnosed with immune-mediated kidney disorders. Additionally, tissue samples were obtained from over 400 patients with clear cell renal cell carcinoma (ccRCC) prior to undergoing immune checkpoint blockade (ICB) therapy. The collected samples were sequenced by multiple platforms such as RNA sequencing (or array), single-cell RNA sequencing (scRNA), or whole exome sequencing (or multi-gene panel sequencing) to generate bulk RNA, scRNA, or DNA data correspondingly. Subsequently, six distinct types of features, referred to as “TIs,” were computed and combined to form the ensemble features. These features encompassed inflammatory expression signatures of genes, pathways, and immune cells, as well as immunogenic mutation signatures of single gene mutations, co-mutations, and tumor mutational burden (TMB). The ensemble of features was utilized to construct an XGBoost machine learning (ML) model. To train and evaluate the performance of the TIs-based ML model for predicting response to ICB therapy and survival outcomes, clinical annotated transcriptome and genome data from about a thousand ccRCC patients were employed.

To investigate the association between inflammatory signatures of autoimmune nephropathy and immunotherapy, we analyzed the expression array data of patients with LN (GSE32591_glo)^36^. In total, we discovered 649 differential pathways and 30 clusters when comparing LN patients to healthy controls for differential expression analysis (Supplementary Fig. 1). Macrophage, and interferon-related pathways were been identified, and those biomarkers have been shown to have a strong correlation with irAE^37^. The results revealed that “regulation of leukocyte mediated immunity” was the most upregulated pathway class in LN, along with “regulation of viral genome replication” and “regulation of defense response to virus by host”, which are related to human endogenous retroviruses (HERVS). The results suggested that ccRCC was an inflammatory tumor, and inflammatory signals were connected to immunotherapy, which was consistent with previous studies^38^.

### Inflammatory signatures were associated with ICB response for autoimmune nephropathy bulk RNA analysis

Inflammation is a complication of autoimmune nephropathy, and inflammatory signatures produced by the immune system are related to the immune response. To investigate the connection between inflammatory signals and immune responses, we disaggregated different types of bulk RNA data from over 1900 patients with immune-mediated kidney disease. After conducting a differential analysis between the cases and healthy controls within each dataset of 5 diseases, we obtained 1,121 differentially expressed genes (DEGs) in LN 55, IgAN 90, MN 586, FSGS 559, and ANCA-AAV 581 separately (Supplementary Fig. 2a). We found that the expression patterns of the top 30 DEGs were generally consistent across distinct datasets, indicating that certain shared molecular processes may contribute to the etiology of diverse autoimmune disorders. (Fig. 2a and Supplementary Data S2). Genes were enriched in the frequent pathways, such as “interferon gamma signaling” pathway, “interferon signaling” pathway, etc., that are relevant to irAE and immune response^39-41^. Among them, the immunoregulation-related gene EGR1 (Early Growth Response 1), involved in Interferon alpha/beta signaling, which is a typical pro-inflammatory cytokine^42^ and enhances anti-tumor capacity by regulating and activating T cells^43^, was differentially expressed in all of the five diseases (Fig. 2a, b). We further evaluated the aforementioned signatures in the IMmotion151 dataset^44^, where 400+ previously untreated ccRCC patients were given atezolizumab (an anti-PD-L1 inhibitor) plus bevacizumab (an anti-VEGF inhibitor), and found 329 genes relevant to response and survival (Fig. 2c). Inflammation is a major class of functions enriched for differentially expressed genes in the autoimmune nephropathy patients compared to healthy controls, and such genes show the potential efficacy to predict response or prognosis of ICB therapy.

**Fig. 2.**
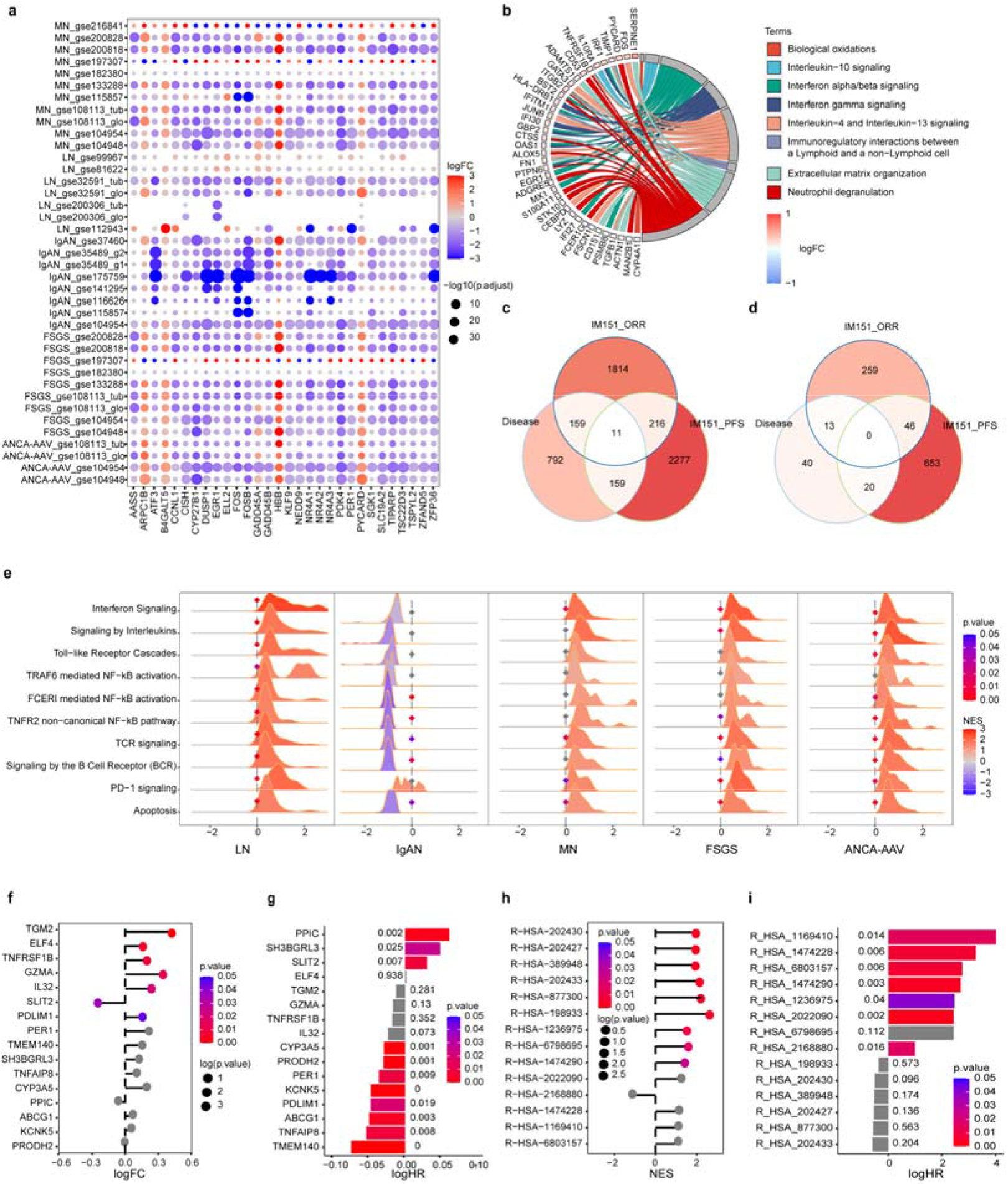
Inflammatory signatures of immune-mediated kidney diseases generated from bulk RNA data. **(a)**The top 30 Differentially Expressed Genes (DEGs) among kidney-mediated diseases between patients and healthy controls in each dataset. The x-axis represents the gene symbols, while the y-axis lists immune-mediated kidney illnesses and the corresponding datasets. Positive logFC represents up-regulated in disease and negative the opposite. **(b)** Representative Reactome pathways and core genes altered in kidney-mediated disease expression in IMmotion151. Positive logFC represents up-regulated in the responders’ group and negative the opposite. **(c-d),** Overlap of signatures from renal disorders and IMmotion151 with those associated with **(c)** genes and **(d)** pathways. **(e)** Differential Expression Pathways (DEPs) in Nephropathy. The ridge maps illustrate the DEPs observed in various types of nephropathies: LN was GSE32591_glo dataset, IgAN was GSE175759 dataset, MN was GSE108113_glo dataset, FSGS was GSE200828 dataset, and ANCA-AAV was GSE108113_glo dataset. **(f-i),** Lollipop charts display kidney disease-derived **(f)** genes and **(h)** pathways expressed differently in responders and non-responders of IMmotion151. Positive logFC represents higher expression in responders and negative the opposite. Bar graphs indicate the correlation of **(g)** gene and **(i)** pathway expression with Progression-Free Survival (PFS) in IMmotion151. Positive logHR represents bad prognosis in higher expression whereas negative logHR represent good prognosis in higher expression.

Furthermore, pathway signatures can replenish the predictions by gene signatures. Within bulk RNA data, we enriched a total of 73 differentially expressed Reactome pathways (DEPs, Supplementary Fig. 2b, 3a and Data S3). Among them, 33 pathways relevant to response and survival in IMmotion 151 (Fig. 2d). In particular, we focused on the canonical pathways most associated with the inflammatory response: interferon pathway; Interleukin pathway; TLR pathway; NF-kB pathway; TCR pathway; BCR pathway, as well as the PD-1 pathway and Apoptosis pathway which are directly related to ICB therapy. The results showed that most of the inflammation-related pathways were significant in at least two or more diseases, especially the TCR pathway and apoptosis pathway were significant in all diseases, and the interleukin pathway and PD-1 pathway were significant in four diseases other than IgAN (Fig. 2e). Intriguingly, most of these signatures up-regulated in responders as compared to non-responders, and prediction ability of pathway signatures were slightly superior to those of gene signatures (Fig. 2f-i). Pathways further explain the changes in inflammatory expression patterns of immune-nephropathy from a functional perspective and can serve as a credible complementary variable in ICB prediction.

Immune cells may be the direct hub of the immunoreactions in both immune diseases and tumors. We employed CIBERSORT to assess the relative proportions of 22 immune cell types in each specimen (Supplementary Fig. 3b). One of the most significantly differentiated cell types, monocytes, was highly concentrated in immune diseases compared to controls. The infiltration and accumulation of them closely related to the pathogenesis of immune diseases^45,46^, and they can also differentiate into dendritic cells to present antigens, or secrete a variety of cytokines to active T cells during tumor immunotherapy^47^ (Supplementary Fig. 3b). In summary, our findings suggest that different levels of inflammatory biomarkers derived from bulk RNA data of immune-mediated kidney diseases can provide valuable insights for predicting ICB outcomes.

### Expansion of inflammatory signatures through single-cell RNA analysis

Single-cell RNA sequencing can effectively complement bulk RNA sequencing through high-resolution expression data. We collected three single-cell RNA datasets (Supplementary Data S1) from LN, IgAN, and MN to perform a differential expression analysis. Specifically, we identified a total of 1,785 DEGs, which included 440, 1264, and 616 genes in LN, IgAN, and MN, respectively (Supplementary Fig. 4a). Compared with bulk RNA data, the scRNA-seq analysis reported 1477 new DEGs, and only 308 (17.3%) DEGs were shared (Supplementary Data S5). This result confirmed that scRNA-seq was capable of complementing signatures at single cell levels. The DEGs identified in scRNA alone were mainly enriched in the “Immune System (HSA-168256)” and “Innate Immune System (HSA-168249)” (Supplementary Data S6). Moreover, “interleukin signaling” pathways associated with genes was upregulated, which turned out to be closely correlated with irAE^37^ and was retrieved by scRNA-seq (Supplementary Fig. 5a). In addition, a total of 518 DEGs were detected that were relevant to response and survival in the IMmotion151 dataset (Fig. 3a). As a result, based on scRNA-seq data, more inflammatory genes involved in immunotherapy may be distinguished.

**Fig. 3.**
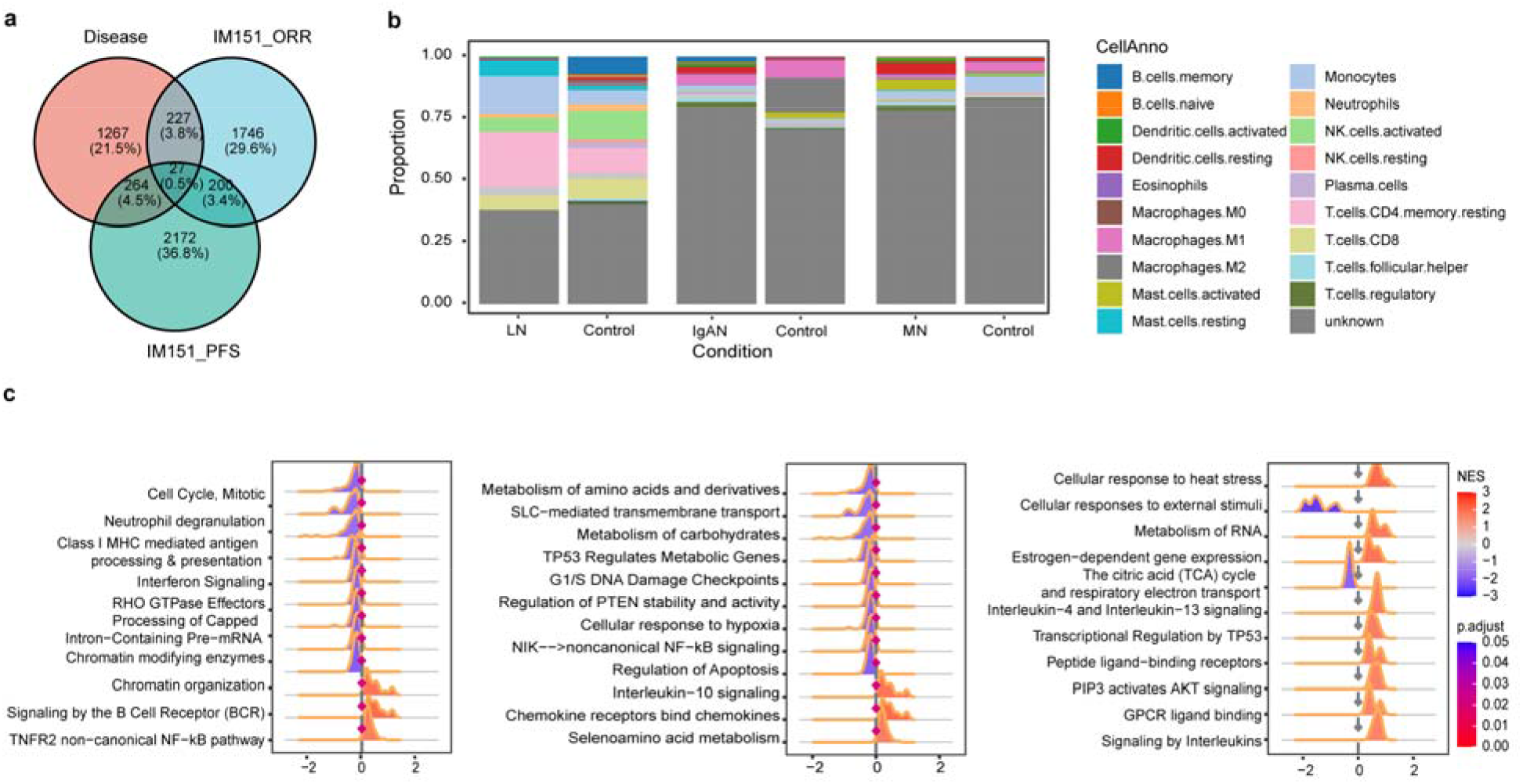
Inflammatory signatures of immune-mediated kidney diseases generated from scRNA data. **(a)**Overlap of gene signatures derived from kidney diseases with those related to response and prognosis in IMmotion151. **(b)** The stacked bar graphs of LN, IgAN, and MN demonstrate the percentage of 22 immune cells in the disease and control groups. **c-e**, the ridge maps represent Differentially Expressed Pathways (DEPs) of the **(c)** LN, **(d)** IgAN, and **(e)** MN datasets, which were obtained from the comparison between patients and healthy controls. Positive normalized enrichment score (NES) represents up-regulated in disease groups and negative the opposite.

Furthermore, the results of DEPs through scRNA-seq data analysis were largely consistent. We additionally discovered 182 DEPs that were associated with immune-mediated kidney diseases, and 145 of these DEPs had inconsistencies with those discovered through bulk RNA datasets (Supplementary Fig. 4b and Supplementary Data S7). DEPs, which were only remembered in scRNA data, were also crucial for immunoregulation and immunoreaction in malignancies (Supplementary Data S7). For example, “DDX58/IFIH1-mediated induction of interferon-alpha/beta (R-HSA-168928)” is an inflammation-related signal but is associated with immune response in tumor immunotherapy based on recognition of intracellular viral RNA and activation of interferon-stimulated gene expression, et al.^48-50^ It was effective in distinguishing immune responders from non-responders (Supplementary Fig. 5b). Above all, 66 pathway signatures were found using scRNA analysis in the IMmotion151 dataset, which was two times the number of those produced from bulk RNA data (Supplementary Fig. 4c). Hence, scRNA-seq data enhanced the accuracy of results by revealing the differences resulting from the immune microenvironment.

Immune cells were annotated by SCINA in the scRNA-seq datasets^51^. We counted the immune cells within the 22 cell types that were associated with kidney diseases using Fisher’s exact test with p.value <0.05 (Fig. 3b and Supplementary Fig. 6). T.cells.CD4.memory.resting was shown to have significantly lower levels in three disease groups compared to healthy controls (Supplementary Fig. 6 and Supplementary Data S8), which was consistent with previous reports^52^.

Eventually, we combined bulk and scRNA signatures by the principle: i) it must become significant in the relevant disease; ii) it in scRNA is needed to be significant generally or in at least two clusters; iii) it will be selected if significant in bulk or single-cell for each disease. As a result, we integrated 1016 genes, 93 pathways, and 16 immune cells of inflammatory signatures from autoimmune nephropathy (Supplementary Fig.11 and Supplementary Data S5-8). In summary, our test emphasized the importance and necessity of adding single-cell RNA data to bulk RNA for inflammatory biomarkers exploring.

### Integrative analysis of inflammatory signatures for predicting immunotherapy response

Obtaining immune-predictive markers in ccRCC patients undergoing immunotherapy may enhance the specificity of immune response predictions. Additionally, the confidence of the indicators can be improved by combining bulk and single-cell RNA data from immunotherapy. Therefore, we mapped the bulk RNA expression data of IMmotion151, divided according to responders (complete response and partial response; CR + PR) and non-responders (stable disease and progressive disease; SD + PD), to the scRNA-Seq data obtained from Bi et al.^53^, which included 4 patients treated with anti-PD1-based therapies (Fig 4a, b, and Supplementary Fig. 7a). This combination approach of bulk and single cells revealed a strong correlation within the responders between the two datasets (Fig. 4c) as well as the two PR patients from Bi et al. (Supplementary Fig. 7b; Pearson correlation = 0.84, p.value = 9.93e-05). Cells in Bi. that had consistent response status (responders or non-responders) with IMmotion151 were kept for further study. Additionally, we got 1711 DEGs and 240 DEPs involving the immune system, hemostasis, extracellular matrix organization, and signal transduction, between mapped effective and ineffective cells. (Fig. 4d, e, and Supplementary Data S5, 7). MT2A, a member of the metallothionein gene family, was obviously upregulated in responders but barely expressed in non-responders, which is consistent with previous findings^54^, hinting at a potential immuno-predictive biomarker (Fig. 4d and Supplementary Fig. 8). We further explored DICs using a similar approach to that used in kidney diseases (Fig. 4f and Supplementary Fig. 9) and tested the prediction ability of these gene and pathway signatures within the IMmotion151 cohort (Supplementary Fig. 10). The aforementioned findings imply that the immune-predictive markers discovered using the mapping approach exhibit a strong correlation with the prognosis of ccRCC patients undergoing immunotherapy.

**Fig. 4.**
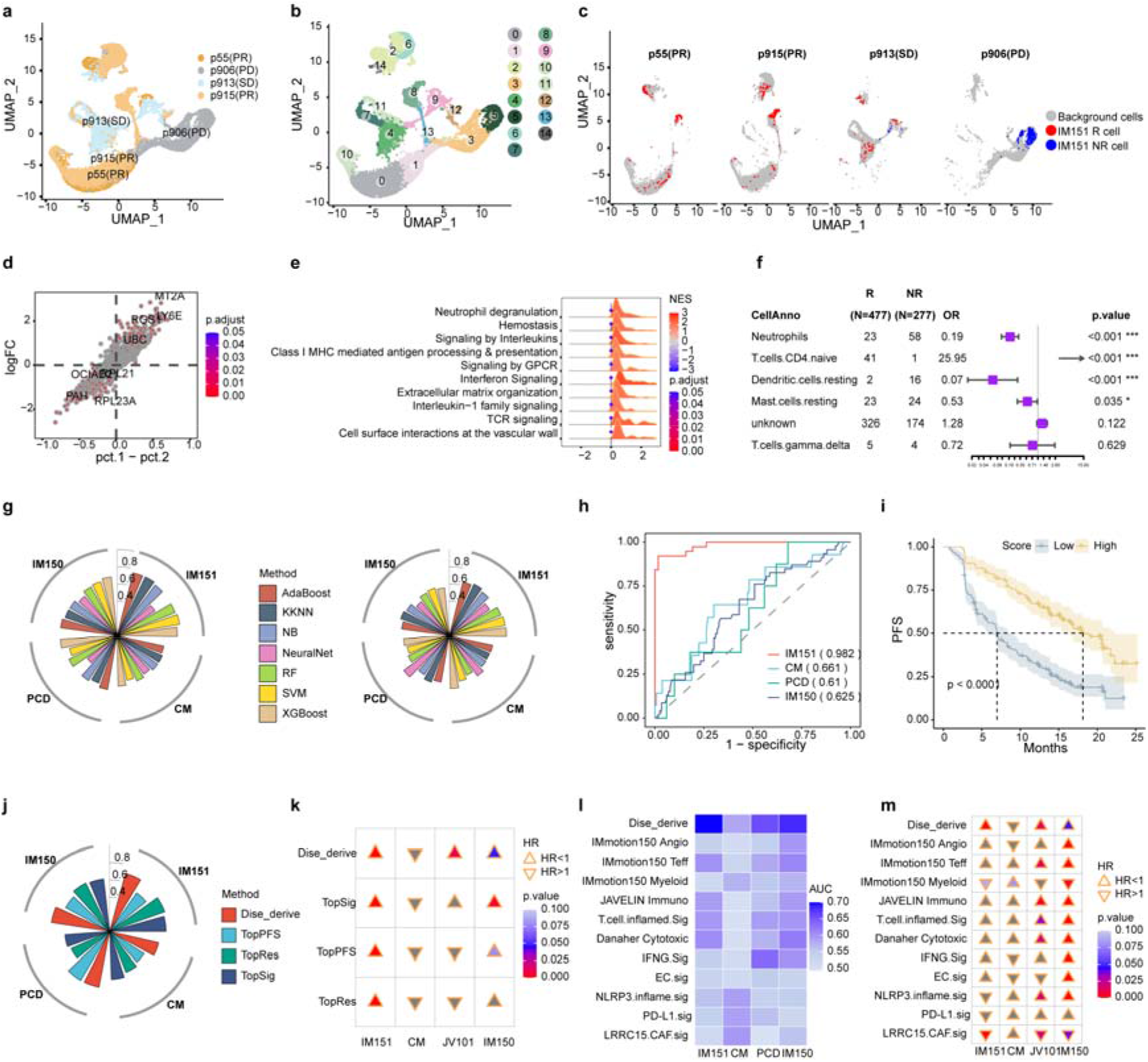
Cell mapping and RNA model generation. **(a-b)**Uniform Manifold Approximation and Projection (UMAP) plots, which show the cell clusters from four patients in Bi et al. **(c)** UMAP plots for the bulk mapping results for each patient. In these plots, responders and non-responders of IMmotion151 mapped to Bi display red dots and blue dots correspondingly. **(d-f)** results of differential analysis from mapped effective and ineffective cells, differentially expressed genes (DEGs, **d**), differentially expressed pathways (DEPs, **e**), and differentially immune cells (DICs, **f**). **(g)** radar maps that compare the AUC of seven machine learning models when gene signatures (left) and pathway signatures (right) from four different ccRCC immune cohorts are used as input. Positive logFC and NES represent higher expression in responders than non-responders and negative the opposite. **(h)** Receiver Operating Characteristic (ROC) curves for the integrated inflammatory features models that evaluated in four ICBs datasets. **(i)** Kaplan-Meier Progress-Free Survival (PFS) curves of inflammatory features models score in IMmotion151. P.value represents the significance of the log-rank test. (**j-k**) efficacy comparisons between model derived from immune-mediated kidney disorders (Dise_derive) and model derived from ICBs. AUC for predicting the response (**j**) and Cox proportional hazards survival test (**h**) in four datasets. The hazard ratio (HR) bigger than 1 in the heatmap represents a bad prognosis for the higher score group. (**l-m**) efficacy comparisons between dise_derive and bio-functional signatures collected from published studies. AUC for predicting the response (**l**) and Cox proportional hazards survival test (**m**) in four datasets.

The mapped DEGs in Bi et al. were intersected with the immune-nephropathy DEGs of bulk and scRNA respectively. The genes common to these two groups were merged, and housekeeping genes were excluded. The protocol of DEP process was the same as DEGs. Finally, 716 genes and 92 pathways specific to immune nephrosis were obtained. Incorporating these markers as intersections with those discovered in the context of renal immune diseases can effectively reduce the number of variables while enhancing the level of confidence. Eventually, 25 genes, 47 pathways, and 4 immune cells were reserved after feature selection by Recursive Feature Elimination (RFE) in the IMmotion151 training datasets (Supplementary Data S9).

Next, we selected seven ML models to compare AUC for multi-genes and multi-pathways individually. XGBoost obtained consistent predictions for response and prognosis by comparing several methods, which was retained for more in-depth study (Fig. 4g and Supplementary Fig. 12). We created an ensemble model using XGBoost based on the models of genes, pathways, and immune cells. And the combined inflammatory signatures ML model could make a robust prediction for ICB response and progression-free survival (PFS) in training data from IMmotion151^55^ (IM151), as well as from four additional independent cohorts: CheckMate^21^ (CM), JAVELIN^18^ (JV101), PCD4989g^56^ (PCD), and IMmotion150^19^ (IM150). The inflammatory ensemble model was able to achieve an AUC of 0.98 in the predictive validation set IMmotion151 (Fig. 4h), and the model’s scores were significantly able to differentiate PFS in IMmotion151 at p.value< 2.18e-20 (Fig. 4i). The performance of the inflammatory model continued to be noteworthy in the independent validation set, with AUCs greater than 0.6 and survival log-rank test significantly in both JAVELIN and IMmotion150 (Fig. 4h and Supplementary Fig. 13b, c). In all datasets, groups with higher scores tended to have a better prognosis than those with lower scores. (Supplementary Fig. 13).

To compare the model efficacy of ccRCC with ICB-derived models and immune-mediated kidney disorders-based models, we build (i) ORR (TopRes); (ii) PFS (TopPFS); (iii) doubly significant genes (TopSig) models (see Method for details). The results indicate that disease-derived gene models perform more effectively and have higher robustness. However, the generalization ability of ICB-derived models’ performance was suboptimal (Fig. 4j and Supplemental Data S13). In terms of prognosis prediction, the disease-derived model discriminates prognosis in three datasets, whereas the best ICB-based model, the doubly significant gene model (TopSig), can only discriminate prognosis in two datasets (Fig. 4k and Supplemental Data S14).

To determine whether diseased-derived models demonstrate superior predictive validity over other biologically sourced models, we compared it to 11 published genetically relevant models (Supplemental Data S11). The results demonstrate that disease-derived models were still the most effective at predicting both response and prognosis, while other models were less effective or could solely predict prognosis but not response (Fig. 4l, m and Supplemental Data S15, S16). For instance, LRRC15.CAF.sig^57^ could predict the prognosis among IMmotion151, JAVELIN, and IMmotion150, but response predictive efficacy AUC was all lower than 0.6. In conclusion, the model based on inflammatory biomarkers is superior to immunotherapy-based signatures models or published features with tumor biology, both in distinguishing immune response and survival for ccRCC with ICB.

### Limitation of immunogenic signatures using genomic data for ICB response assessment

Immunogenic biomarkers are critical in assessing the immune response. However, the efficacy of using a single biomarker to predict the response to immunotherapy in ccRCC is constrained. The calculation of TMB, which includes all nonsynonymous mutations, demonstrated modest predictive capacity with an AUC of 0.572 in the IMmotion151 cohort (Supplementary Fig. 18). Using Fisher’s exact test and Cox proportional hazards regression analysis, we identified genes that were differentially mutated in response or prognosis with ICB. In the end, 14 mutation genes were selected as potential immunogenic signatures (Fig. 5a, b, and Supplementary Data S10).

**Fig. 5.**
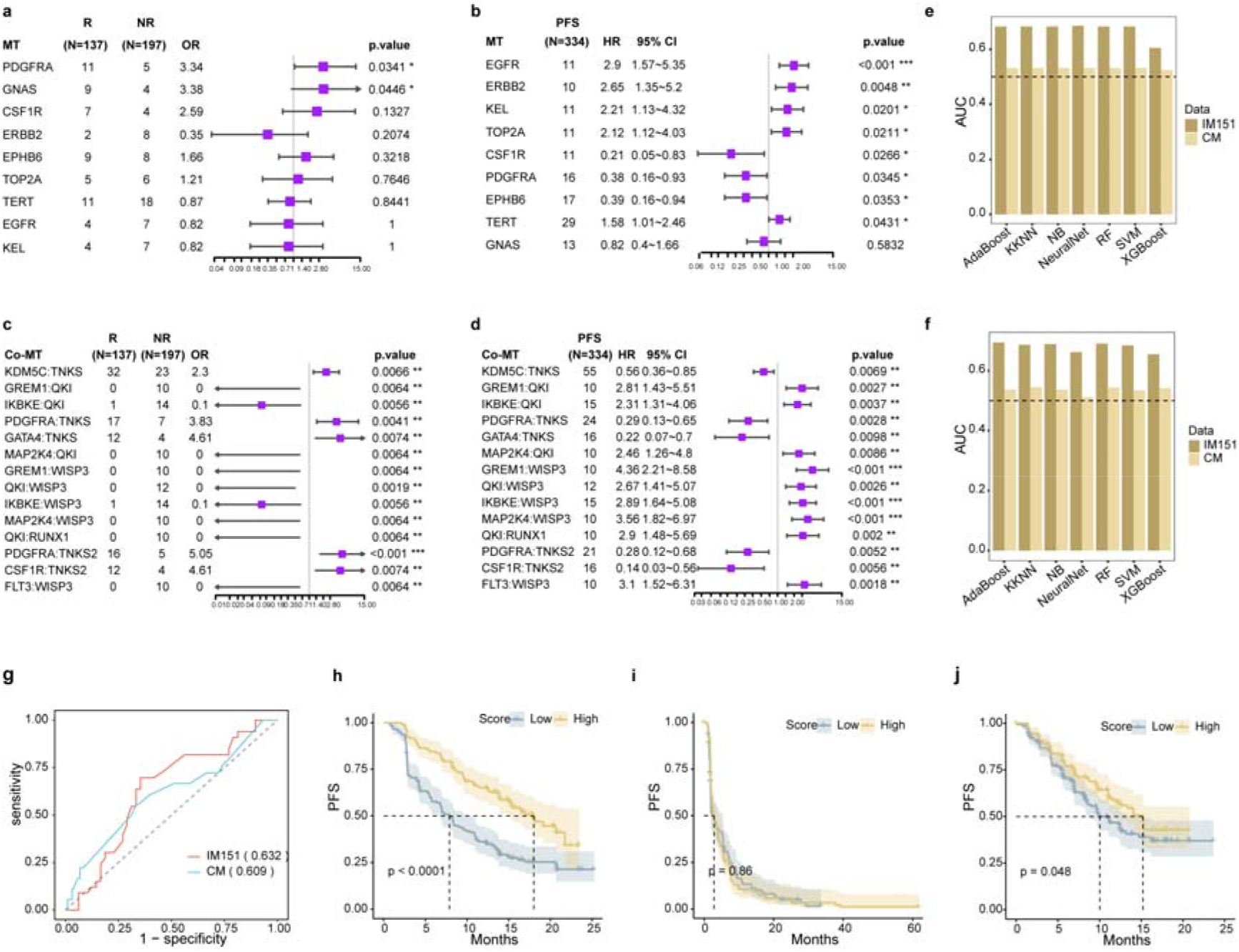
Immunogenic Signatures-Based Predictive Model for Immunotherapy Outcomes. **a-d,**depict the predictive values of single gene mutations for **(a)** response and **(b)** survival outcomes, as well as co-mutations for **(c)** response and **(d)** survival outcomes. (**e-f),** AUC comparison of seven distinct machine learning models, with the result of single gene mutations and **(f)** co-mutations in immunological cohorts. **(g)** ROC curves for the integrated immunogenic signature models that were evaluated in the two DNA data available ICBs datasets. **(h-j)** Kaplan-Meier survival curves of immunogenic signatures models score in IMmotion151 (**h**) CheckMate (**i**) and JAVELIN (**j**).

Notably, the highest population frequency of the concurrent mutation gene pairs, EPHB6-GNAS, only in 3 patients (Supplementary Fig. 14). We calculated co-mutation genes, which are pairs of genes that either one of them were mutated. Hence, we screened for co-mutated gene pairs in the IMmotion151 cohort by analyzing the differences in immune response and prognosis. In total, we identified 1876 combinations of gene pairs (Fig. 5c, d and Supplementary Data S10). To reduce selection bias, we filtered additional co-mutation genes based on the overall survival of The Cancer Genome Atlas (TCGA) in KIRC. Ultimately, we discovered 49 co-mutation genes associated with response and survival in patients.

We compared seven different approaches to building machine learning models based on multigene and co-mutation genes. XGBoost consistently demonstrated satisfactory performance in predicting response and prognosis using the IMmotion151 or CheckMate cohorts (Fig. 5e,f and Supplementary Fig. 15-16). Overall, predictive models based on the three individual types of immunogenic signatures have demonstrated limited ability to estimate immune response and survival (Fig. 5e,f and Supplementary Fig. 15-16). Finally, we established a machine-learning model that integrates all the immunogenic biomarkers. The model achieved an AUC of 0.632 and 0.609 in the testing set IMmotion151 and the independent validation set CheckMate cohort, respectively (Fig. 5g). The comprehensive model’s predictive score was capable of stratifying the prognosis of different cohorts (Fig. 5h-j). These results revealed that the single immunogenic model was inadequate, and the multi-indicator combination model improved the predictive efficiency of immunotherapy.

### Integration of TIs-based machine learning tests for ICB effectiveness prediction

The inflammatory model was more efficient than the immunogenic model, but there’s still room for improvement in the inflammatory model. Two models derived from the distinct perspective and integration of the two may further enhance the predictive efficiency. We aimed to develop a machine learning model using multi-omics data to predict response in ccRCC patients receiving ICB treatment. The multi-omics signatures landscape was shown in Fig. 6a and Supplementary Fig. 17. The ML-based multi-omics model was constructed using 354 patients with ICB from IMmotion151 who had RNA-seq and DNA data. The patients were randomly assigned into a training and validation set with a ratio of 2:1. Our TIs-ML model achieved an excellent AUC of 0.997 for distinguishing responders from non-responders in the validation set, significantly surpassing the predictive power of the GEP score (AUC: 0.627), TME score (AUC: 0.574), TIDE score (AUC: 0.560), TMB (AUC: 0.572), and PD-L1 expression (AUC: 0.555) (Fig. 6b and Supplementary Fig. 18b). Positive findings were consistently observed when the model was applied to the CheckMate cohort; our model had an AUC of 0.705, whereas the GEP score had 0.498, the TME score had 0.509, and the TIDE score had 0.573 (Fig. 6c). The present study suggests that the TIs-based predictive signatures are highly generalizable and that the TIs-ML model can robustly predict response to ICB.

**Fig. 6.**
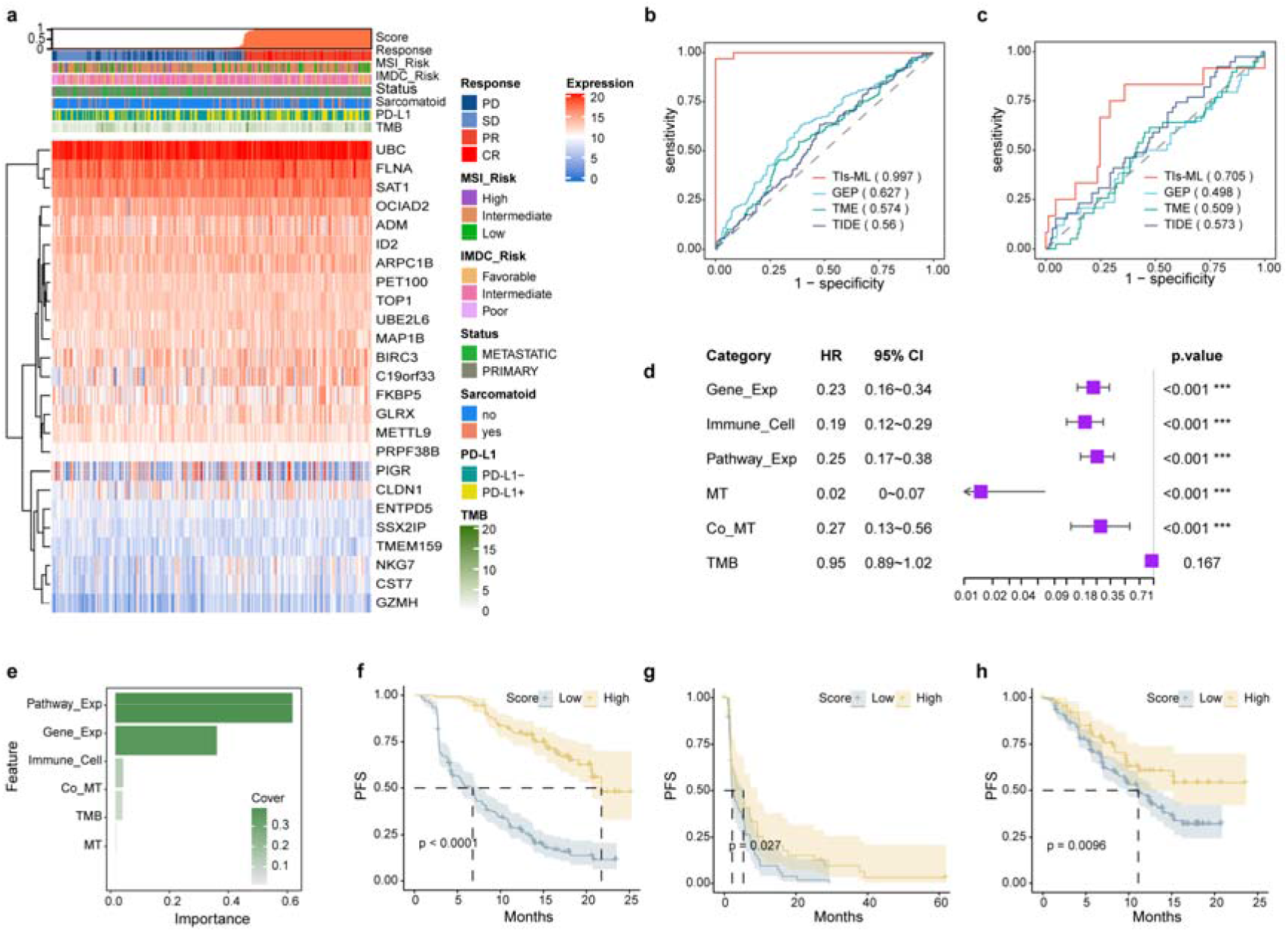
Multi-Omics model for predicting immunotherapy outcomes. **(a)**Expression profiles of 25 model gene signatures in the IMmotion151 dataset are depicted, representing a comprehensive view of the landscape. (**b-c),** ROC curves of response prediction efficacy comparison between multi-omics (TIs) model and previous immune-related ICBs prediction model in IMmotion151 (b) and CheckMate (c). **(d)** Forest map of prognostical efficacy of each TIs signature for survival in IMmotion151 **(e)** The importance coefficient of each signature in contributing to the overall mode is illustrated. (**f-h),** Kaplan-Meier curves in the cohorts of **(f)** IMmotion151, **(g)** CheckMate, and **(h)** JAVELIN demonstrate the effectiveness of the TIs approach in predicting PFS.

In addition, individuals in the high-score group (with the threshold set at the median) in the IMmotion151 cohort had a longer PFS than those in the low-score group (Figure 6f; p.value < 0.0001). The CheckMate and JAVELIN101 cohorts (Fig. 6g,h and Supplementary Fig.19) showed a better prognosis in the high-score group (p.value = 0.027 for PFS and 0.54 for overall survival in the CheckMate cohort; p.value = 0.010 in the JAVELIN101 cohort). Similarly, The TIs-ML score outperformed typical biomarkers for predicting prognosis (Supplementary Fig.21). These findings demonstrated that the TIs score could act as a prognostic biomarker for better patient stratification in ccRCC with ICB treatment.

More importantly, we revealed that inflammatory signatures demonstrated a higher importance coefficient than immunogenic signatures (Fig. 6e) in the multi-omics model. Autoimmune nephropathy-related pathways and genes played a dominant role in the model (Fig. 6d), suggesting that there was a strong correlation between immune-mediated kidney disease and immunotherapy.

### Association of risk score with clinical factors and biological characteristics

We examined the correlation between the TIs-ML scores and clinical factors after observing a significant connection between the risk score and tumor regression (Fig. 7a). A significant difference in TIs score was observed between sarcomatoid and non-sarcomatoid ccRCC, with sarcomatoid ccRCC tending to have a higher TIs score (Fig. 7e, f). Our study also identified that the Memorial Sloan-Kettering Cancer Center (MSKCC)^58^ and the International Metastatic Renal Cell Carcinoma Database Consortium (IMDC)^59,60^, which are commonly used to predict the prognosis in metastatic RCC using serological biomarkers, were less effective compared with TIs score (Supplementary Fig. 23a-b, d-e). The results demonstrated that there was no significant difference in TIs score between primary and metastatic lesions (Supplementary Fig. 23c, f), which emphasizes the robustness of our model in predicting response to ICB.

**Fig. 7.**
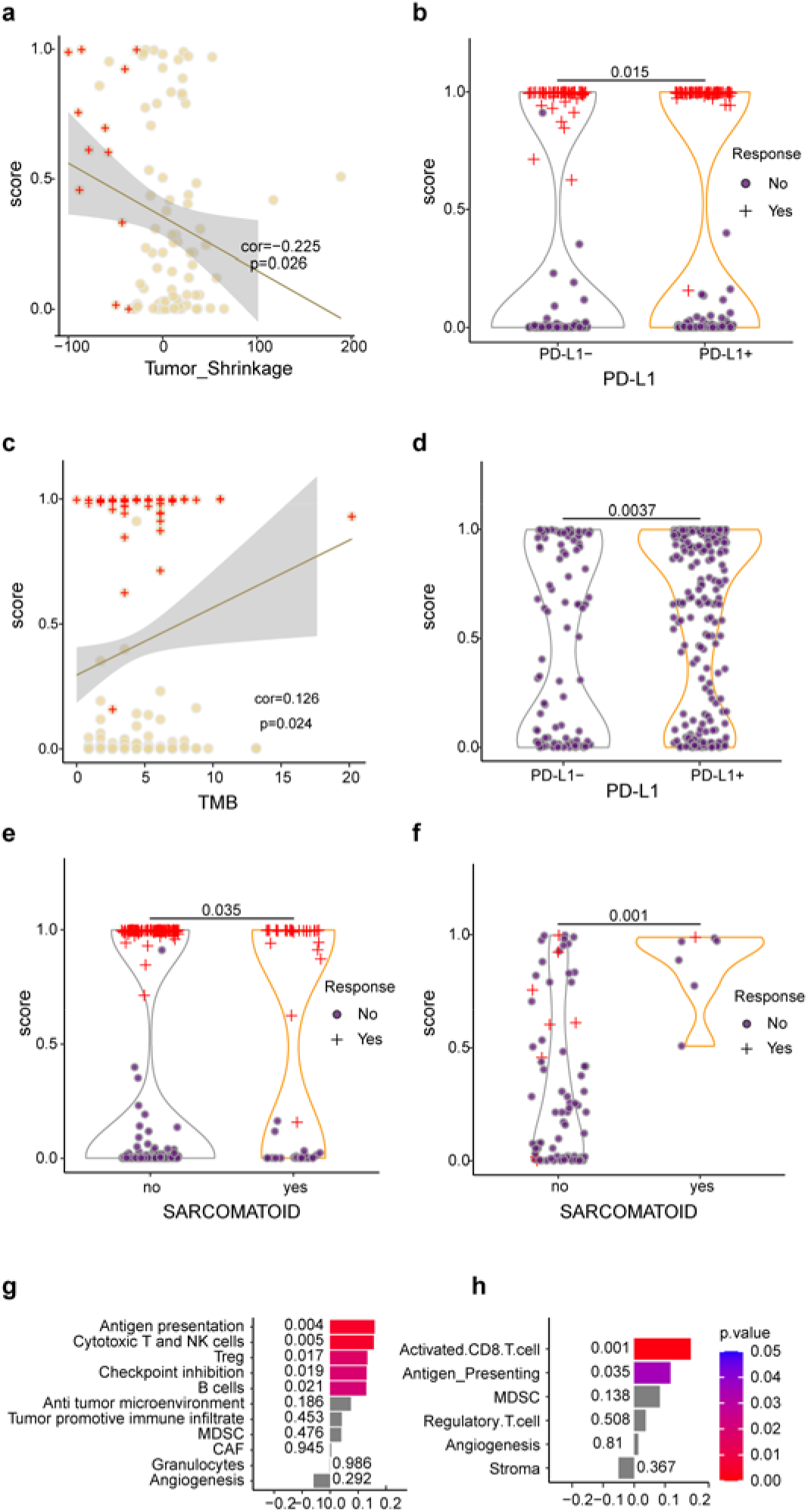
Correlations between the model score and various biological and clinicopathologic factors. The model score shows correlations with **(a)** tumor regression and **(c)** tumor mutation burden (TMB) in the IMmotion151 dataset. In addition, the model score is correlated with PD-L1 expression in both the **(b)** IMmotion151 and **(d)** JAVELIN datasets. Furthermore, the model score exhibits correlations with sarcomatoid differentiation in the **(e)** IMmotion151 and **(f)** CheckMates datasets. Moreover, the model score is correlated with tumor microenvironment-related signatures derived from **(g)** Bagaev.sig.2 and **(h)** Multi_Source.sig. The X-axis represents the correlations between TIs score and TME-related signatures.

Furthermore, we explored the correlation between TIs scores and well-known immunotherapy biomarkers, including TMB and PD-L1. Responders were more closely related to high scores than high TMB (Fig. 7c) and positive PD-L1(Fig. 7b,d). Besides, immunotherapy responders and survival-benefit patients with TMB-low or PD-L1-negative could be bailed by higher TIs-based risk scores, emphasizing that our multi-omics model can complement the prediction ability of conventional biomarkers (Supplementary Fig. 24).

We examined the association between risk scores and published ICB-related signatures, and identified a positive correlation between the risk scores and a variety of immune-related hallmark genes (Supplementary Data S12). The inflammatory response enhances the efficiency of immune checkpoint therapy by increasing immune cell infiltration and improving tumor immunogenicity (Supplementary Fig. 25a and Supplementary Fig. 26). The study showed that the TIs score had a positive correlation with activated CD8 T cells and antigen-presenting signatures, as well as novel gene expression signatures such as NK cells, Treg, and B cells in the analysis of the TME (Fig. 7g, h and Supplementary Figure 25b, Supplementary Data S12). These immune cell signatures are known to play a key role in the tumor-killing process.

### TIs-ML was specific for ccRCC patients treated with immunotherapy

Since the model score can predict the response and prognosis of immunotherapy in ccRCC, we tested the specificity of our model by evaluating non-immunotherapy treatment for ccRCC, and also by assessing immunotherapy treatment for other cancer patients. The risk score was ineffective in predicting response or prognosis between the single-agent targeted therapy and antiangiogenic therapy in ccRCC from the IMmotion151, JAVELIN, and CheckMate cohorts (Fig. 8a-d). The IMvigor210 (mUC)^61^ and Poplar (NSCLC)^62^ datasets were also analyzed using the ML-based inflammatory model with negative results for predicting prognosis or response (Fig. 8e-h and Supplementary Fig. 27a, b). In conclusion, our method can specifically predict ICB outcomes in ccRCC patients but is not suitable for predicting efficacy in ccRCC patients receiving other therapies or for individuals with other types of cancer undergoing immunotherapy.

**Fig. 8.**
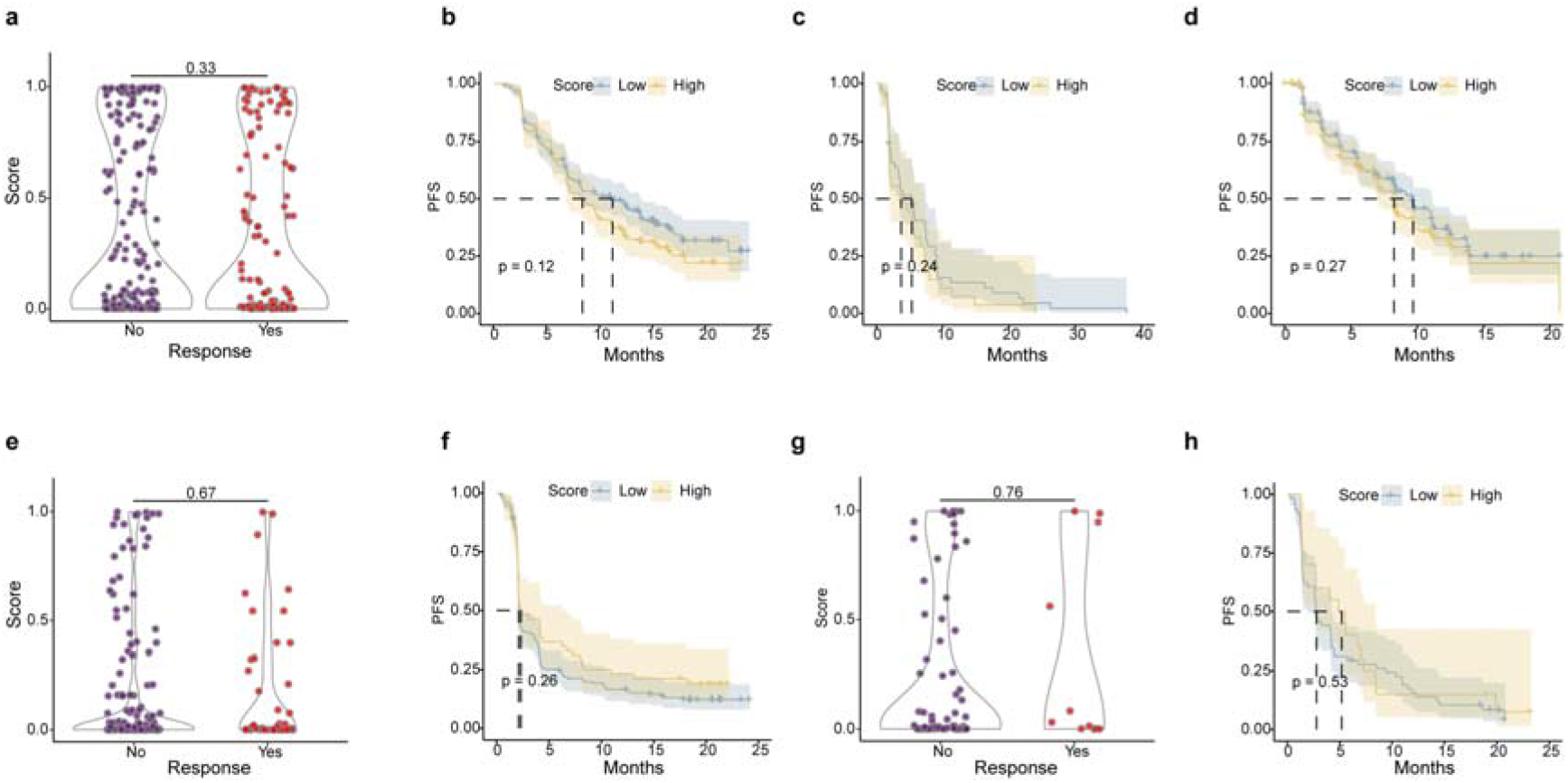
The multi-omics model is specifically designed for ccRCC patients undergoing immunotherapy. Our model cannot predict the responses of individuals treated with **(a)** sunitinib in IMmotion151 (ccRCC), **(e)** atezolizumab in IMvigor210 (mUC), and **(g)** atezolizumab in Poplar (NSCLC). Additionally, our model cannot predict the progression-free survival (PFS) of patients treated with **(b)** sunitinib in IMmotion151 (ccRCC), **(c)** everolimus in CheckMate (ccRCC), **(d)** sunitinib in JAVELIN (ccRCC), atezolizumab in IMvigor210 (mUC), and **(h)** atezolizumab in Poplar (NSCLC).

### Inflammation-associated differential expression gene in autoimmune nephropathy differentiating the result of immunotherapy

To further explore the roles and mechanisms of signatures derived from immune-mediated kidney disorders in guidance ICBs for ccRCC, biological function pathway enrichment analysis was performed for the disease-derived genes.

Pathway enrichment analysis indicated that immune-mediated kidney disease genes have a strong association with biological processes, which play a crucial role in immune stress and evasion in tumors (Supplementary Fig.29, Fig. 2e, and Fig. 3c). Furthermore, we performed a consensus clustering analysis using these genes in the IMmotion151 dataset and identified two distinct clusters (Fig. 9a). C1 and C2 differentially expressed genes were mainly enriched in seven pathways, including “regulation of lymphocyte activation” and “nuclear division”, which play a significant role in immunogenic cell death and exhibit high expression levels in C1 compared to C2 (Fig. 9b). Interestingly, patients in Cluster 1 had a poorer prognosis. However, Immunotherapy improved PFS in the C1 patients while in the C2, neither immunotherapy nor any other therapy affected the prognosis (Fig. 9d and Supplementary Fig.30, 31). Consensus cluster analysis suggests that cluster1 patients are more suitable for monotherapy or combination therapy of ICB, while cluster2 could select more economical therapy in ccRCC. However, immune kidney disease signatures are likely only applicable to KIRC and may not be generalizable to most other cancer types in pan-cancer analysis (Fig. 9c, e, f and Supplementary Fig.32, 33). In summary, the results demonstrate that kidney disease-related genes have a very strong correlation with immunotherapy at the molecular level and can specifically predict immunotherapy in renal cancer.

**Fig. 9.**
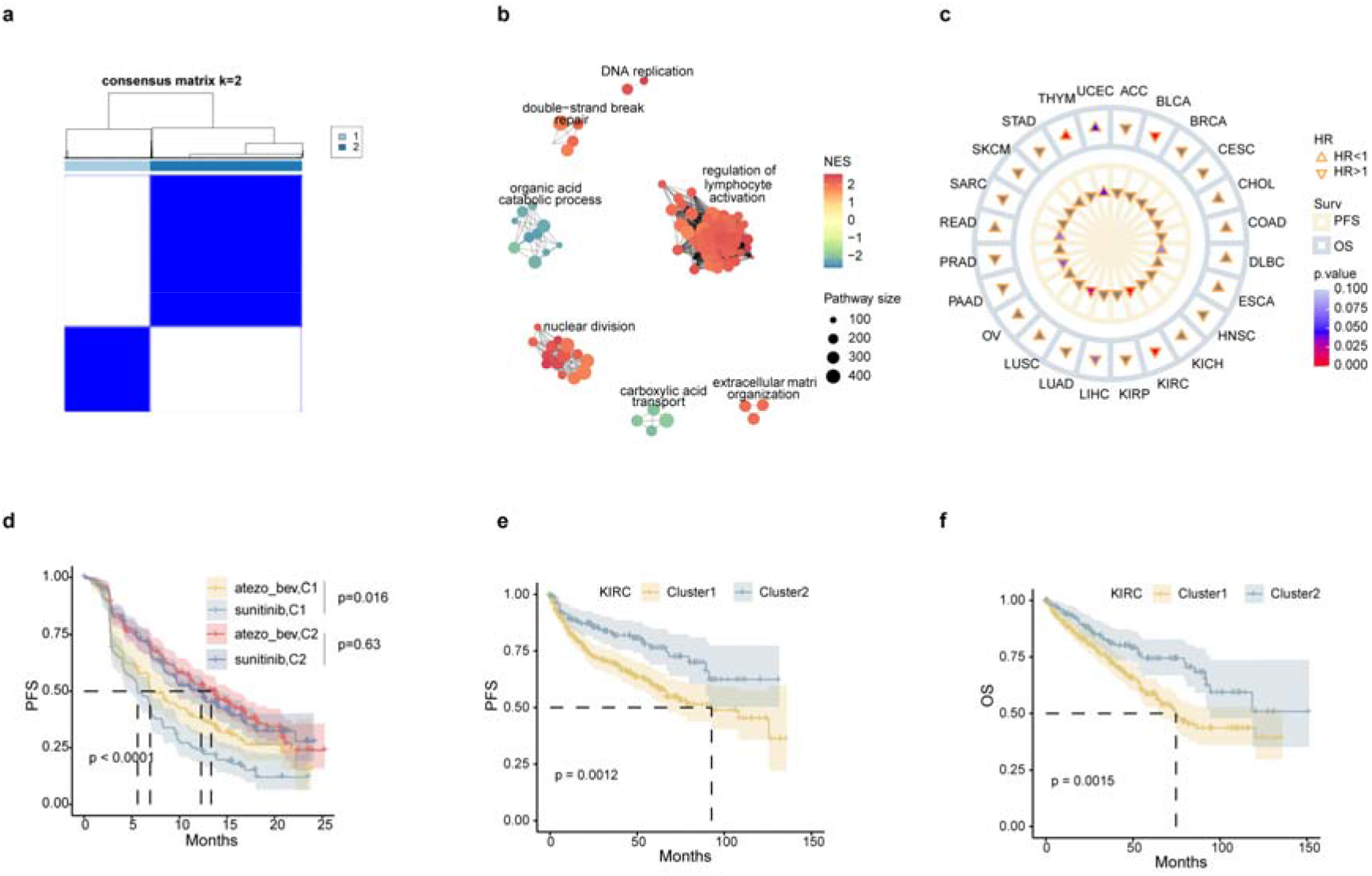
Consensus clustering of biomarkers obtained from immunological diseases. **(a)**Concensus clustering correlation heatmaps of IMmotion151 samples that genes associated with nephropathy were subjected to two consensus clusters. **(b)** Net graph of GO biological process pathways which significantly altered in Cluster 1 versus Cluster 2. Positive normalized enrichment score (NES) represents up-regulated in Cluster1, and Negative NES represents down-regulated in Cluster1. (**c**) Kaplan-Meier curves of the combination of treatment and clusters that demonstrated ICBs were superior to target monotherapy in cluster1. **(d)** clusters prognostical effect on PFS and overall survival (OS) in TCGA pan-cancer cohorts. HR>1 represents that Cluster1 was the bad prognosis indicator compare with Cluster2 and HR<1 is the opposite. Clusters were significant in stratifying PFS (**e**) and OS (**f**) in TCGA KIRC.

## Discussion

In this study, we aimed to develop an immunotherapy predictive approach for all patients with ccRCC that is accurate, reliable, and precise. Our model is superior to the well-established biomarkers like TMB and PD-L1. Our model accurately and robustly predicted the response of ICB therapy, regardless of whether it was anti-PD1 or PD-L1 therapy, first-line or second-line therapy, or monotherapy or combination therapy.

We speculate that there may be connections between immune diseases and immunotherapy in tumors because cancer can suppress or escape the immune reaction and immunoreaction triggered by anti-PD-1 axis inhibitors may cause autoimmune diseases^63^. A preliminary experiment in LN vs normal analysis demonstrated that immune kidney disease correlated with up-regulated expression of “regulation of leukocyte mediated immunity” and pathways related to HERVS. Pathways enrichment clusters could correspond to the factors related to kidney ICB therapy. Besides, we explained this hypothesis by consensus clustering analysis for nephropathy-related genes resulting in two clusters indicating distinct prognoses. Specifically, ccRCC patients in cluster 1 had poor prognoses but were more likely to benefit from combination programs of anti-PD-(L)1 plus multi-TKI (sunitinib). We were surprised that cluster 1 might not be a good indicator of immune monotherapy (Fig. 9a, b), with a possible mechanism for sunitinib can enhance tumor infiltration by downregulating immunosuppressive cytokines and depleting Treg cells and MDCSs in murine models^64^. Our results suggest that the exploration of immune-predictive indicators from renal disease is on the right track.

Increasingly more and more studies demonstrate that multi-omics features have advantages in accurately predicting outcomes. For example, Vanguri et al. integrated radiology, pathology, and genomics to predict ICB response in non-small cell lung cancer, reaching an AUC of 0.8, which outperformed unimodal of TMB (AUC=0.61) and PD-L1 (AUC=0.73)^65^. Newell et al. combined genomics, methylomics, transcriptomics, and immune cell infiltrates to predict ICB response in melanoma with an AUC of 0.84^66^. In our test, we incorporated bulk RNA profiles, single-cell RNA profiles, and genomics to build the model to predict ICB response and prognosis in ccRCC, achieving excellent prediction efficacy across multiple cohorts.

However, our study has certain limits. All the cohorts employed were public datasets with a limited number of patients, and there was a lack of independent validation datasets from prospective clinical cohorts. In the next phase, we plan to recruit patients for the proposed immunotherapy, regardless of the number of lines of therapy or the treatment regimen. Multi-omics assays are both expensive and time-consuming for clinical use, making them a limitation. Alternative approaches are necessary to modify the variables to better align with clinical practice. Additionally, gene expression values are known to be less stable and can vary significantly among platforms, batches, and standardized forms. To address this issue, we conducted a study that translated gene expression values into the order of a gene’s expression level in a single sample. Preliminary results are available and will be published soon.

## Methods

### Data collection

We systematically searched the transcriptome data of 5 immune-mediated kidney diseases from 36 datasets in the Gene Expression Omnibus (GEO) database^67^ based on the review articles^34,35^. These datasets included bulk RNA datasets of 5 LN datasets (GSE112943, GSE200306, GSE81622, GSE81622, GSE99967), 7 IgAN datasets (GSE175759, GSE37460, GSE104954, GSE141295, GSE116626, GSE35489, GSE115857), 10 MN datasets (GSE216841, GSE200828, GSE200818, GSE182380, GSE197307, GSE133288, GSE115857, GSE108113, GSE104954, GSE104948), 8 FSGS datasets (GSE200828, GSE200818, GSE182380, GSE197307, GSE133288, GSE108113, GSE104954, GSE104948) and 3 ANCA-AAV datasets (GSE108113, GSE104954, GSE104948), as well as 3 scRNA datasets of IgAN (GSE171314), MN (GSE171458), and LN (Arazi et al.^68^) (Supplementary Data S1). It should be noted that we excluded cases and controls that were less than 5 in bulk datasets and those with obviously non-active LN in GSE99967. We also excluded one IgAN patient with microproteinuria in GSE17131.

For the immune datasets, we collected transcriptome and genomic data with clinical annotations from three ccRCC cohorts: IMmotion151^44^ (atezolizumab + bevacizumab versus Sunitinib; first-line) from the European Genome-phenome Archive (EGA) dataset, CheckMate (nivolumab versus everolimus; latter-line) from the supplementary materials provided by Braun et al^21^, and JAVELIN (avelumab + axitinib versus sunitinib; first-line) from the supplementary materials provided by Motzer et al.^18^.We also gathered 4 clinically annotated transcriptome immune cohorts of urothelial carcinoma (UC) (IMvigor210^61^, treated by atezolizumab, n = 208), NSCLC (POPLAR^62^, treated by atezolizumab, n = 81), RCC (IMmotion150^19^, treated by atezolizumab alone or in combination with bevacizumab, n = 162), multi-cancer (PCD4989g^56^, treated by atezolizumab, n = 206). Patients with clearly 4 treatment outcomes of “CR, PR, PD, SD” were enrolled except JEVELIN which with known survival time records were enrolled.

Mutation and clinical data of ccRCC from The Cancer Genome Atlas (TCGA) database were downloaded from cbioportal (https://www.cbioportal.org/study/summary?id=kirc_tcga). This study generates no new data, so no ethics approval is needed. The Supplemental Data S1 of this study contains comprehensive information on all of the data.

### Bulk RNA-seq data process and analysis

All expression array data were log2-transformed using the GEO protocol. RNA-seq count data were normalized to FPKM for each gene based on the gencode v22 and log2-transformed, subsequently. The analysis of DEGs was performed using limma (R package, v3.2.3), with significant differences in gene expression in a single dataset being determined by p.value<0.05 and abs(logFC)>1. Disease-associated genes were required to satisfy the requirement of being significant in three or more datasets, with the ANCA-AAV threshold set at two for only four datasets, exceptionally.

### scRNA-seq data process and analysis

In the single-cell dataset, data were processed and analyzed using Seurat (R package, v3.2.3), and batch effect correction was performed using harmony (R package, v0.1.1). Significance thresholds in the single-cell analysis were set at p<0.05 and avg_logFC>0.25. The differential genes between immune-mediated kidney disorders and health from a single-cell perspective were determined to be significant either in the holistic condition or in more than two clusters.

### Analysis of bulk RNA-seq mapped to scRNA-seq

Only four samples in the Bi dataset had information on the known response of ICB treatment, and the samples treated with atezolizumab + bevacizumab in IMmotion151 were mapped to the Bi dataset using Scissor (R package, v2.0.0) corresponding to the parameter’s alpha=0.05 and cutoff=0.25. For the results of the mapping only the cells with consistent response were retained, i.e., those whose outcome was the same as either response or non-response in the Bi. and in IMmotion151 cells were included in the downstream analysis. The mapped cells were still analyzed for DEGs between the response vs. non-response groups using Seruat, and significant DEGs were identified using p.value<0.05 and avg_logFC>0.25.

### Pathway and immune cell annotation and selection

All significant genes in each dataset were enriched pathways using the enrichPathway function from ReactomePA (R package, v1.30) and the profiling of 22 immune cells computed by CIBERSORT (Bulk RNA-seq) and SCINA (scRNA-seq), respectively. Autoimmune nephropathy-associated pathways need to meet the above criteria consistent with immune-nephropathy genes. Differential immune cells were screened in bulk using the Mann-Whitney U test and in scRNA-seq using the Fisher’ test. Immune cells associated with kidney disease also needed to meet the following criteria: (i) significant in three or more bulk datasets; (ii) significant in single-cell conditions; and (iii) significant in either bulk or scRNA for the same disease.

### ICB Data Preprocessing

The log2FPKM matrices of IMmotion151, IMmotion150, JAVELIN, and PCD4989g were individually 0.75 quantile normalized, and combined with the CheckMate data, followed by batch correction using the ComBat function in the SVA (R package, v3.34). The expression matrix of genes for pre-feature selection was a subset of autoimmune nephropathy-related genes. Pathway expression matrix analysis was performed using the ssGESA (GSVA, R package, v1.34) and subset to immune-mediated kidney disease pathways.

For the mutation matrix of all ICB datasets, only non-synonymous mutations were taken into account in this study. Candidate single-mutation variables that were significantly related to response (Fisher’s exact exact test, p.value<0.05) and survival (Cox-proportional hazards regression model, Wald test, p.value<0.05) in IMmotion151. Co-mutation genes are defined as pairs of genes that have either mutated or not. We filtered candidate co-mutation variables by response or survival in IMmotion151 and retained only those that were significantly associated with overall survival with KIRC in TCGA.

### Feature selection and modeling

Five model variables need feature selection ulteriorly: genes expression, pathways expression, immune cell proportion, genes mutation, and co-mutation. Recursive feature reduction using caret (R package, v6.0-93) with 10-fold cross-validation was performed on all candidate variables. ML models were created using a variety of algorithms, including support vector machine (SVM), Naïve Bayes (NB), random forest (RF), k-nearest neighbors (KKNN), AdaBoost Classification Trees (AdaBoost), eXtreme Gradient Boosting (XGBoost), Neural Network (NeuralNet). The caret package (R package, v1.7.5.1) provides the first five methods, nerualnet package (R package, v1.44.2) provides the NeuralNet algorithm, and XGBoost obtained from the xgboost package (R package, v6.0-93).

### Comparison of different multi-gene models

To evaluate the performance of the models, we compared them with the ICB-derived gene model and biological functional characterization models sequentially. For ICB-derived genes selected by: (i) top 25 response significant genes (TopRes); (ii) top 25 survival significant genes (TopPFS); (iii) combined top 25 significant genes in response and survival (survival and response multiplied by p.value and sort in ascending order then took the top 25 genes, TopSig). The ICB-derived model built by XGBoost was applicable to the TIs-ML model in the procedure. The set of ICB-related functional gene lists were collected from multiple studies and is detailed in Supplementary Data S12. Expression scores for the above functional signatures were calculated by the ssGSEA function from GSVA (R package, v1.34), and the pROC (R package, v1.18) was used to generate the ROC curves by calculating the sensitivity and specificity of the scores in predicting the ICBs’ binary responses. Other ICB response prediction model included that: the GEP score was computed by 18 immune genes expression which formula provided in Keynote-028^69^; the TME score was computed by TMEscore (R package, v0.1.4); TIDE score was computed by the TIDEpy python package (https://github.com/liulab-dfci/TIDEpy).

### Identifying clusters of autoimmune-nephropathy associated genes

716 genes were identified in the immune-mediated kidney disorders samples from the above analytic processes and excluded the housekeeping genes. We clustered the expression matrix of immune-nephropathy genes of IMmotion151 using ConsensusClusterPlus (R package, v1.50), resulting in two distinct consensus that cluster by k-means and distance computed by Euclidean. The cosine similarity was determined by calculating the top 50 genes that were specifically highly expressed in each cluster. For the assignment of the CheckMate and JAVELIN sample clusters, greater cosine similarity with the known IMmotion151 clusters was labeled corresponding clusters (a similarity lower than 0.4 would be assign in NA).

### Statistical analysis

R v3.6.3 (https://www.r-project.org) was used to perform all statistical analyses. The two-sided Mann-Whitney U test was used to compare the difference in scores between the response and non-response (as well as other clinical) groups. Fisher’s exact test was utilized to determine whether there is a significant correlation between discrete variables and clinical groups. The logarithmic ranking test was used to assess prognostic differences between subgroups of clinical or model scores for Kaplan-Meier (KM) survival curves. Adjusted p.value of DEGs and GSEA analysis computed by the Benjamini-Hochberg method. By default, continuous variables greater than the median or quantile of 0.75 would be classified as high. The correlation between model score and immunological characteristics or cancer hallmarks was evaluated using Spearman correlation.

## Data available

All the data mentioned in this paper are public data, and detailed descriptions are given in the Methods section. Algorithms used for data analysis are all publicly available from the indicated references in the Methods section.

## Abbreviation

Tis: inflammatory and immunogenic signatures
ccRCC: clear cell renal cell carcinoma
ICBs: immune checkpoint blockades
PD-1: programmed cell death 1
PD-L1: programmed death ligand-1
CTLA-4: cytotoxic T lymphocyte antigen 4
TMB: tumor mutational burden
DEGs: differentially expressed genes
DEPs: differentially expressed pathways
DICs: differential immune cells
PFS: progression-free survival
OS: overall survival
TME: tumor immune microenvironment
RNA-seq: bulk RNA sequencing
scRNA-seq: single-cell RNA sequencing
ML: machine learning
irAE: immune-related adverse effects
LN: lupus nephritis
IgAN: IgA nephropathy
MN: membranous nephropathy
FSGS: focal segmental glomerulosclerosis
ANCA-AAV: antineutrophil cytoplasmic antibody vasculitis
HERVS: human endogenous retroviruses

## References

1. Maio, M., et al. Pembrolizumab in microsatellite instability high or mismatch repair deficient cancers: updated analysis from the phase II KEYNOTE-158 study. Ann Oncol 33, 929–938 (2022).

2. Shitara, K., et al. Nivolumab plus chemotherapy or ipilimumab in gastro-oesophageal cancer. Nature 603, 942–948 (2022).

3. Motzer, R.J., et al. Nivolumab versus Everolimus in Advanced Renal-Cell Carcinoma. N Engl J Med 373, 1803–1813 (2015).

4. Motzer, R.J., et al. Avelumab plus Axitinib versus Sunitinib for Advanced Renal-Cell Carcinoma. N Engl J Med 380, 1103–1115 (2019).

5. Rini, B.I., et al. Pembrolizumab plus Axitinib versus Sunitinib for Advanced Renal-Cell Carcinoma. N Engl J Med 380, 1116–1127 (2019).

6. Choueiri, T.K., et al. Nivolumab plus Cabozantinib versus Sunitinib for Advanced Renal-Cell Carcinoma. N Engl J Med 384, 829–841 (2021).

7. Lee, C.H., et al. Lenvatinib plus pembrolizumab in patients with either treatment-naive or previously treated metastatic renal cell carcinoma (Study 111/KEYNOTE-146): a phase 1b/2 study. Lancet Oncol 22, 946–958 (2021).

8. Rini, B.I., et al. Atezolizumab plus bevacizumab versus sunitinib in patients with previously untreated metastatic renal cell carcinoma (IMmotion151): a multicentre, open-label, phase 3, randomised controlled trial. Lancet 393, 2404–2415 (2019).

9. Motzer, R.J., et al. Nivolumab plus Ipilimumab versus Sunitinib in Advanced Renal-Cell Carcinoma. N Engl J Med 378, 1277–1290 (2018).

10. Reck, M., et al. Pembrolizumab versus Chemotherapy for PD-L1-Positive Non-Small-Cell Lung Cancer. N Engl J Med 375, 1823–1833 (2016).

11. Hodi, F.S., et al. Nivolumab plus ipilimumab or nivolumab alone versus ipilimumab alone in advanced melanoma (CheckMate 067): 4-year outcomes of a multicentre, randomised, phase 3 trial. Lancet Oncol 19, 1480–1492 (2018).

12. Hsu, C., et al. Safety and Antitumor Activity of Pembrolizumab in Patients With Programmed Death-Ligand 1-Positive Nasopharyngeal Carcinoma: Results of the KEYNOTE-028 Study. J Clin Oncol 35, 4050–4056 (2017).

13. Rizvi, N.A., et al. Cancer immunology. Mutational landscape determines sensitivity to PD-1 blockade in non-small cell lung cancer. Science 348, 124–128 (2015).

14. Samstein, R.M., et al. Tumor mutational load predicts survival after immunotherapy across multiple cancer types. Nat Genet 51, 202–206 (2019).

15. Marabelle, A., et al. Association of tumour mutational burden with outcomes in patients with advanced solid tumours treated with pembrolizumab: prospective biomarker analysis of the multicohort, open-label, phase 2 KEYNOTE-158 study. Lancet Oncol 21, 1353–1365 (2020).

16. Meiri, E., et al. Pembrolizumab (P) in patients (Pts) with colorectal cancer (CRC) with high tumor mutational burden (HTMB): Results from the Targeted Agent and Profiling Utilization Registry (TAPUR) Study. Journal of Clinical Oncology 38, 133–133 (2020).

17. Labriola, M.K., et al. Characterization of tumor mutation burden, PD-L1 and DNA repair genes to assess relationship to immune checkpoint inhibitors response in metastatic renal cell carcinoma. J Immunother Cancer 8(2020).

18. Motzer, R.J., et al. Avelumab plus axitinib versus sunitinib in advanced renal cell carcinoma: biomarker analysis of the phase 3 JAVELIN Renal 101 trial. Nat Med 26, 1733–1741 (2020).

19. McDermott, D.F., et al. Clinical activity and molecular correlates of response to atezolizumab alone or in combination with bevacizumab versus sunitinib in renal cell carcinoma. Nat Med 24, 749–757 (2018).

20. Krishna, C., et al. Single-cell sequencing links multiregional immune landscapes and tissue-resident T cells in ccRCC to tumor topology and therapy efficacy. Cancer Cell 39, 662–677 e666 (2021).

21. Braun, D.A., et al. Interplay of somatic alterations and immune infiltration modulates response to PD-1 blockade in advanced clear cell renal cell carcinoma. Nat Med 26, 909–918 (2020).

22. Motzer, R.J., et al. Molecular Subsets in Renal Cancer Determine Outcome to Checkpoint and Angiogenesis Blockade. Cancer Cell 38, 803–817 e804 (2020).

23. Sammut, S.J., et al. Multi-omic machine learning predictor of breast cancer therapy response. Nature 601, 623–629 (2022).

24. Liu, Z., et al. Integrated multi-omics profiling yields a clinically relevant molecular classification for esophageal squamous cell carcinoma. Cancer Cell 41, 181–195 e189 (2023).

25. Kong, J., et al. Network-based machine learning approach to predict immunotherapy response in cancer patients. Nat Commun 13, 3703 (2022).

26. Lech, M. & Anders, H.J. The pathogenesis of lupus nephritis. J Am Soc Nephrol 24, 1357–1366 (2013).

27. He, J.W., Zhou, X.J., Lv, J.C. & Zhang, H. Perspectives on how mucosal immune responses, infections and gut microbiome shape IgA nephropathy and future therapies. Theranostics 10, 11462–11478 (2020).

28. Grandi, N. & Tramontano, E. HERV Envelope Proteins: Physiological Role and Pathogenic Potential in Cancer and Autoimmunity. Front Microbiol 9, 462 (2018).

29. Masetti, R., et al. Autoimmunity and cancer. Autoimmun Rev 20, 102882 (2021).

30. Puzanov, I., et al. Managing toxicities associated with immune checkpoint inhibitors: consensus recommendations from the Society for Immunotherapy of Cancer (SITC) Toxicity Management Working Group. J Immunother Cancer 5, 95 (2017).

31. Haanen, J., et al. Management of toxicities from immunotherapy: ESMO Clinical Practice Guidelines for diagnosis, treatment and follow-up. Ann Oncol 28, iv119-iv142 (2017).

32. Socinski, M.A., et al. Association of Immune-Related Adverse Events With Efficacy of Atezolizumab in Patients With Non-Small Cell Lung Cancer: Pooled Analyses of the Phase 3 IMpower130, IMpower132, and IMpower150 Randomized Clinical Trials. JAMA Oncol 9, 527–535 (2023).

33. van Not, O.J., et al. Association of Immune-Related Adverse Event Management With Survival in Patients With Advanced Melanoma. JAMA Oncol 8, 1794–1801 (2022).

34. Kant, S., Kronbichler, A., Sharma, P. & Geetha, D. Advances in Understanding of Pathogenesis and Treatment of Immune-Mediated Kidney Disease: A Review. Am J Kidney Dis 79, 582–600 (2022).

35. Boesen, E.I. & Kakalij, R.M. Autoimmune-mediated renal disease and hypertension. Clin Sci (Lond) 135, 2165–2196 (2021).

36. Berthier, C.C., et al. Cross-species transcriptional network analysis defines shared inflammatory responses in murine and human lupus nephritis. J Immunol 189, 988–1001 (2012).

37. Jing, Y., et al. Multi-omics prediction of immune-related adverse events during checkpoint immunotherapy. Nat Commun 11, 4946 (2020).

38. Hegde, P.S. & Chen, D.S. Top 10 Challenges in Cancer Immunotherapy. Immunity 52, 17–35 (2020).

39. Hu, W.T., et al. Eosinophil and IFN-gamma associated with immune-related adverse events as prognostic markers in patients with non-small cell lung cancer treated with immunotherapy. Front Immunol 14, 1112409 (2023).

40. Head, L., et al. Biomarkers to predict immune-related adverse events with checkpoint inhibitors. Journal of Clinical Oncology 37, 131–131 (2019).

41. Ayers, M., et al. IFN-gamma-related mRNA profile predicts clinical response to PD-1 blockade. J Clin Invest 127, 2930–2940 (2017).

42. Pozhitkov, A.E., et al. Tracing the dynamics of gene transcripts after organismal death. Open Biol 7(2017).

43. Dan, L., et al. The phosphatase PAC1 acts as a T cell suppressor and attenuates host antitumor immunity. Nat Immunol 21, 287–297 (2020).

44. Buttner, F.A., et al. A novel molecular signature identifies mixed subtypes in renal cell carcinoma with poor prognosis and independent response to immunotherapy. Genome Med 14, 105 (2022).

45. Zheng, Y., et al. Single-Cell Transcriptomics Reveal Immune Mechanisms of the Onset and Progression of IgA Nephropathy. Cell Rep 33, 108525 (2020).

46. Tang, R., et al. A Partial Picture of the Single-Cell Transcriptomics of Human IgA Nephropathy. Front Immunol 12, 645988 (2021).

47. Ugel, S., Cane, S., De Sanctis, F. & Bronte, V. Monocytes in the Tumor Microenvironment. Annu Rev Pathol 16, 93–122 (2021).

48. Yoneyama, M. & Fujita, T. Function of RIG-I-like receptors in antiviral innate immunity. J Biol Chem 282, 15315–15318 (2007).

49. Loo, Y.M., et al. Distinct RIG-I and MDA5 signaling by RNA viruses in innate immunity. J Virol 82, 335–345 (2008).

50. Honda, K., Yanai, H., Takaoka, A. & Taniguchi, T. Regulation of the type I IFN induction: a current view. Int Immunol 17, 1367–1378 (2005).

51. Zhang, Z., et al. SCINA: A Semi-Supervised Subtyping Algorithm of Single Cells and Bulk Samples. Genes (Basel) 10(2019).

52. Okoye, A.A. & Picker, L.J. CD4(+) T-cell depletion in HIV infection: mechanisms of immunological failure. Immunol Rev 254, 54–64 (2013).

53. Bi, K., et al. Tumor and immune reprogramming during immunotherapy in advanced renal cell carcinoma. Cancer Cell 39, 649–661 e645 (2021).

54. Deng, T., et al. Single cell sequencing revealed the mechanism of PD-1 resistance affected by the expression profile of peripheral blood immune cells in ESCC. Front Immunol 13, 1004345 (2022).

55. Motzer, R.J., et al. Final Overall Survival and Molecular Analysis in IMmotion151, a Phase 3 Trial Comparing Atezolizumab Plus Bevacizumab vs Sunitinib in Patients With Previously Untreated Metastatic Renal Cell Carcinoma. JAMA Oncol 8, 275–280 (2022).

56. Herbst, R.S., et al. Predictive correlates of response to the anti-PD-L1 antibody MPDL3280A in cancer patients. Nature 515, 563–567 (2014).

57. Dominguez, C.X., et al. Single-Cell RNA Sequencing Reveals Stromal Evolution into LRRC15(+) Myofibroblasts as a Determinant of Patient Response to Cancer Immunotherapy. Cancer Discov 10, 232–253 (2020).

58. Motzer, R.J., et al. Survival and prognostic stratification of 670 patients with advanced renal cell carcinoma. J Clin Oncol 17, 2530–2540 (1999).

59. Heng, D.Y., et al. Prognostic factors for overall survival in patients with metastatic renal cell carcinoma treated with vascular endothelial growth factor-targeted agents: results from a large, multicenter study. J Clin Oncol 27, 5794–5799 (2009).

60. Ko, J.J., et al. The International Metastatic Renal Cell Carcinoma Database Consortium model as a prognostic tool in patients with metastatic renal cell carcinoma previously treated with first-line targeted therapy: a population-based study. Lancet Oncol 16, 293–300 (2015).

61. Balar, A.V., et al. Atezolizumab as first-line treatment in cisplatin-ineligible patients with locally advanced and metastatic urothelial carcinoma: a single-arm, multicentre, phase 2 trial. Lancet 389, 67–76 (2017).

62. Fehrenbacher, L., et al. Atezolizumab versus docetaxel for patients with previously treated non-small-cell lung cancer (POPLAR): a multicentre, open-label, phase 2 randomised controlled trial. Lancet 387, 1837–1846 (2016).

63. Yasunaga, M. Antibody therapeutics and immunoregulation in cancer and autoimmune disease. Semin Cancer Biol 64, 1–12 (2020).

64. Petroni, G., Buque, A., Zitvogel, L., Kroemer, G. & Galluzzi, L. Immunomodulation by targeted anticancer agents. Cancer Cell 39, 310–345 (2021).

65. Vanguri, R.S., et al. Multimodal integration of radiology, pathology and genomics for prediction of response to PD-(L)1 blockade in patients with non-small cell lung cancer. Nat Cancer 3, 1151–1164 (2022).

66. Newell, F., et al. Multiomic profiling of checkpoint inhibitor-treated melanoma: Identifying predictors of response and resistance, and markers of biological discordance. Cancer Cell 40, 88–102 e107 (2022).

67. Edgar, R., Domrachev, M. & Lash, A.E. Gene Expression Omnibus: NCBI gene expression and hybridization array data repository. Nucleic Acids Res 30, 207–210 (2002).

68. Arazi, A., et al. The immune cell landscape in kidneys of patients with lupus nephritis. Nat Immunol 20, 902–914 (2019).

69. Ott, P.A., et al. T-Cell-Inflamed Gene-Expression Profile, Programmed Death Ligand 1 Expression, and Tumor Mutational Burden Predict Efficacy in Patients Treated With Pembrolizumab Across 20 Cancers: KEYNOTE-028. J Clin Oncol 37, 318–327 (2019).

