## supplementary_figure for "Multi-omics features-based machine learning method improve immunotherapy response in clear cell renal cell carcinoma"

### Slide 1
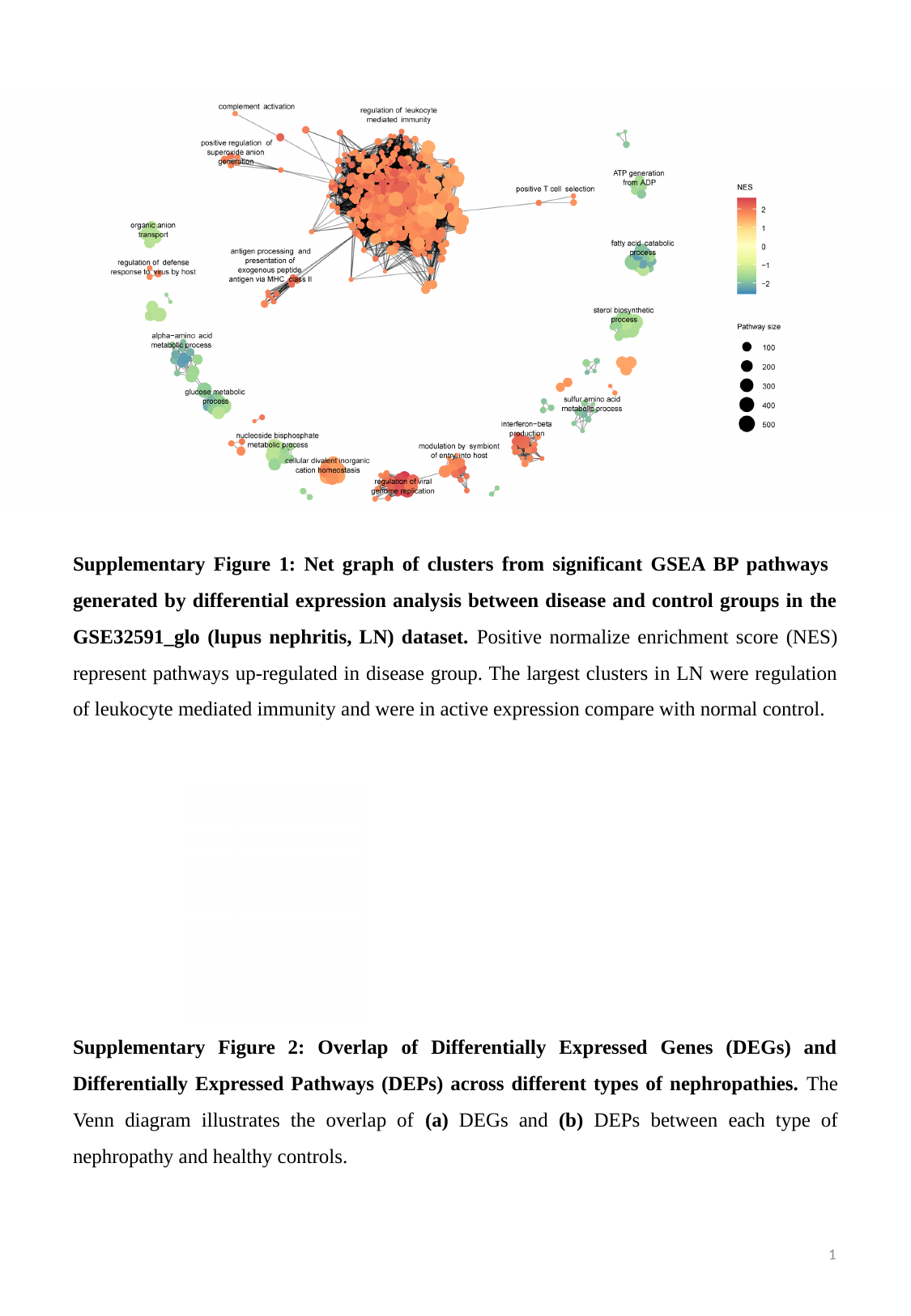

Supplementary Figure 1: Net graph of clusters from significant GSEA BP pathways generated by differential expression analysis between disease and control groups in the GSE32591_glo (lupus nephritis, LN) dataset. Positive normalize enrichment score (NES) represent pathways up-regulated in disease group. The largest clusters in LN were regulation of leukocyte mediated immunity and were in active expression compare with normal control.
Supplementary Figure 2: Overlap of Differentially Expressed Genes (DEGs) and Differentially Expressed Pathways (DEPs) across different types of nephropathies. The Venn diagram illustrates the overlap of (a) DEGs and (b) DEPs between each type of nephropathy and healthy controls.
1

### Slide 2
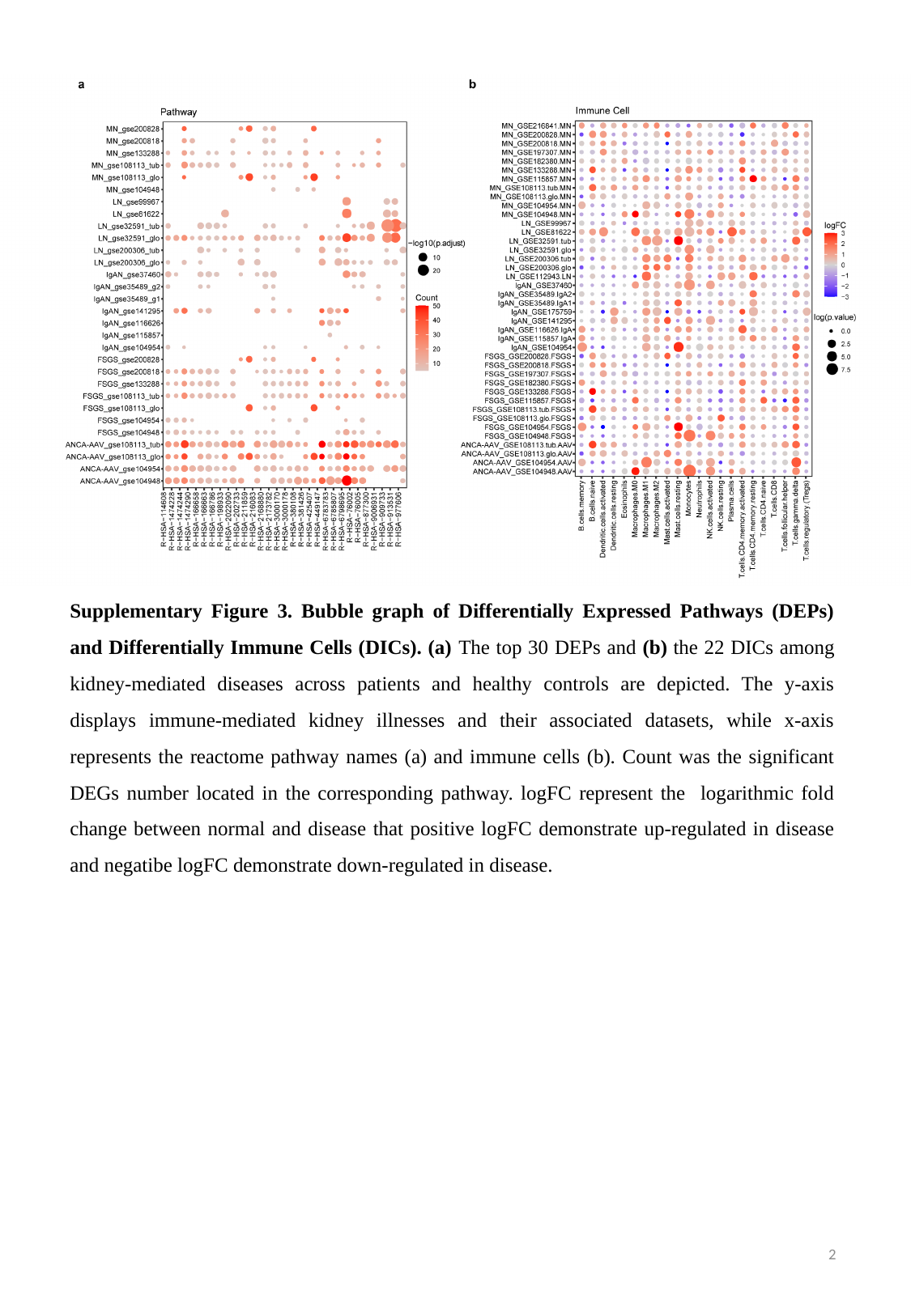

Supplementary Figure 3. Bubble graph of Differentially Expressed Pathways (DEPs) and Differentially Immune Cells (DICs). (a) The top 30 DEPs and (b) the 22 DICs among kidney-mediated diseases across patients and healthy controls are depicted. The y-axis displays immune-mediated kidney illnesses and their associated datasets, while x-axis represents the reactome pathway names (a) and immune cells (b). Count was the significant DEGs number located in the corresponding pathway. logFC represent the logarithmic fold change between normal and disease that positive logFC demonstrate up-regulated in disease and negatibe logFC demonstrate down-regulated in disease.
2

### Slide 3
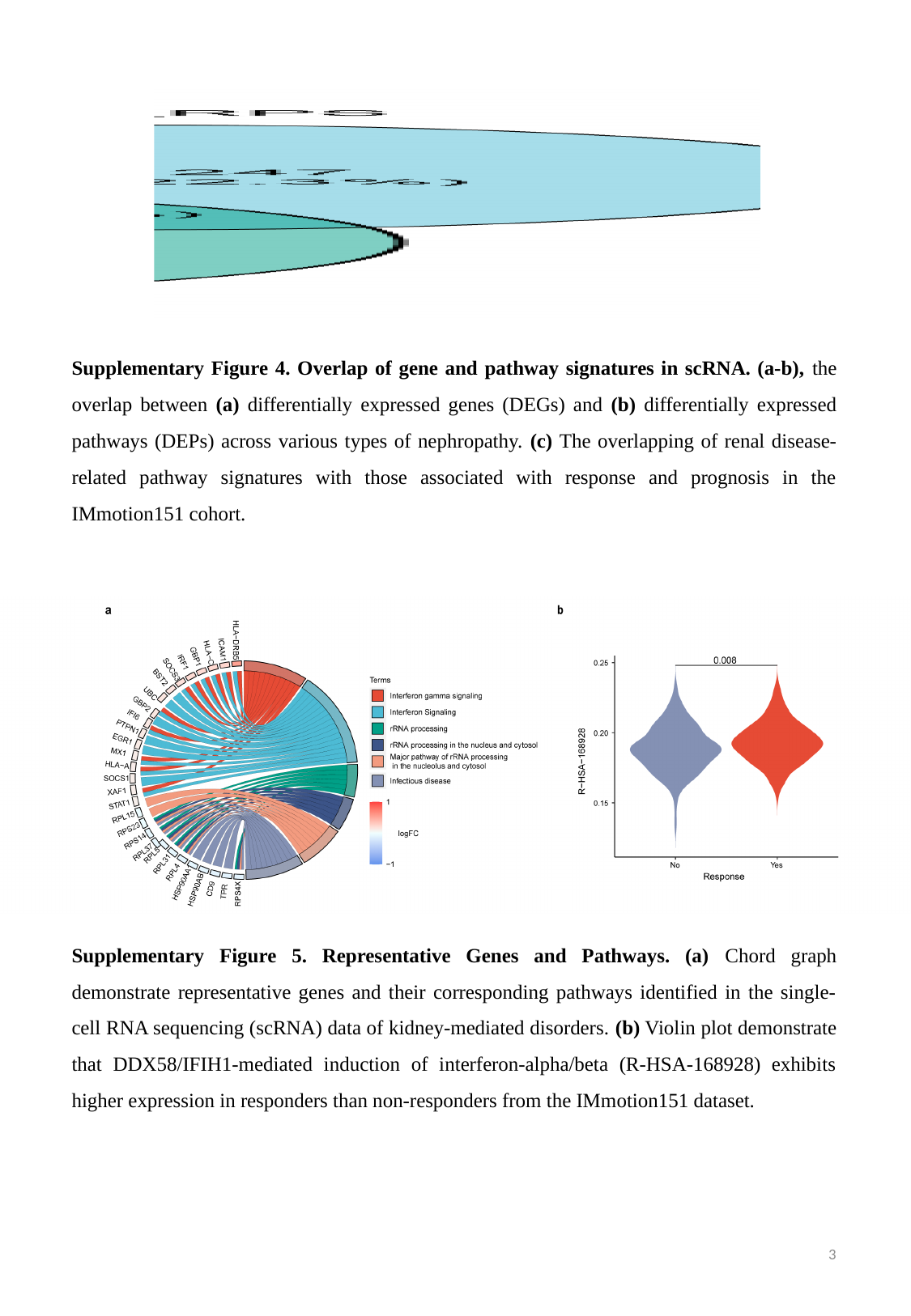

Supplementary Figure 4. Overlap of gene and pathway signatures in scRNA. (a-b), the overlap between (a) differentially expressed genes (DEGs) and (b) differentially expressed pathways (DEPs) across various types of nephropathy. (c) The overlapping of renal disease-related pathway signatures with those associated with response and prognosis in the IMmotion151 cohort.
Supplementary Figure 5. Representative Genes and Pathways. (a) Chord graph demonstrate representative genes and their corresponding pathways identified in the single-cell RNA sequencing (scRNA) data of kidney-mediated disorders. (b) Violin plot demonstrate that DDX58/IFIH1-mediated induction of interferon-alpha/beta (R-HSA-168928) exhibits higher expression in responders than non-responders from the IMmotion151 dataset.
3

### Slide 4
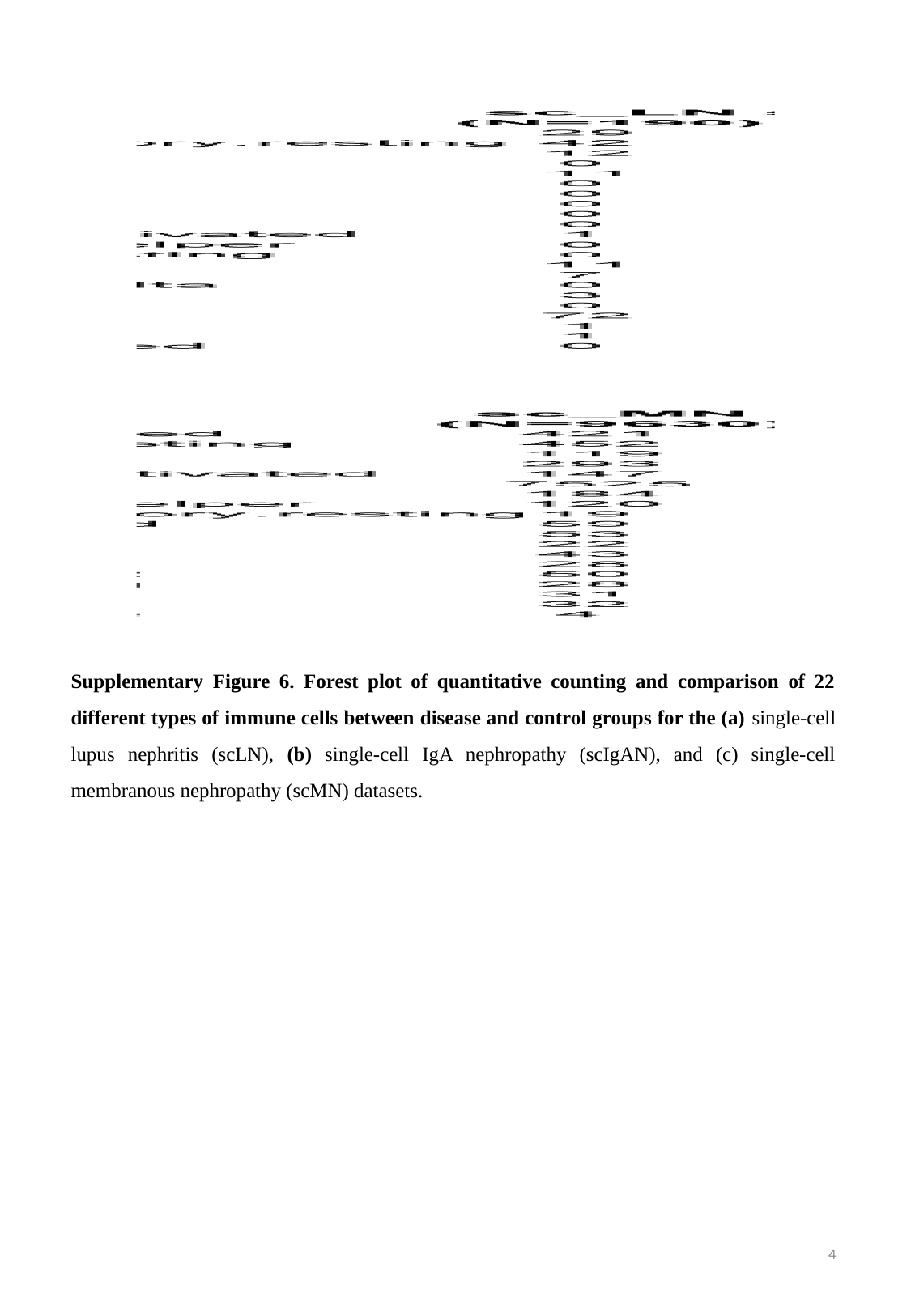

Supplementary Figure 6. Forest plot of quantitative counting and comparison of 22 different types of immune cells between disease and control groups for the (a) single-cell lupus nephritis (scLN), (b) single-cell IgA nephropathy (scIgAN), and (c) single-cell membranous nephropathy (scMN) datasets.
4

### Slide 5
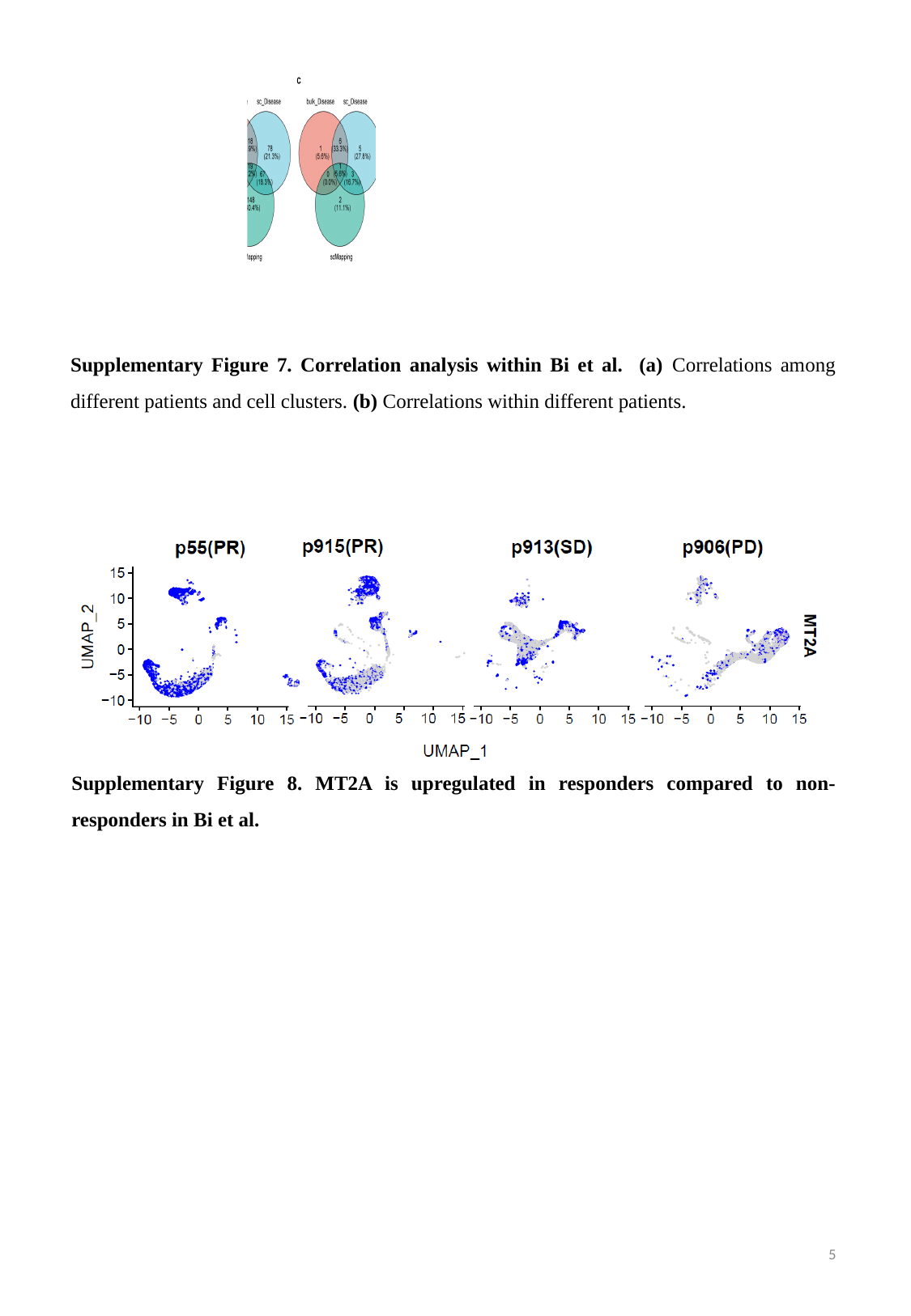

Supplementary Figure 7. Correlation analysis within Bi et al. (a) Correlations among different patients and cell clusters. (b) Correlations within different patients.
Supplementary Figure 8. MT2A is upregulated in responders compared to non-responders in Bi et al.
5

### Slide 6
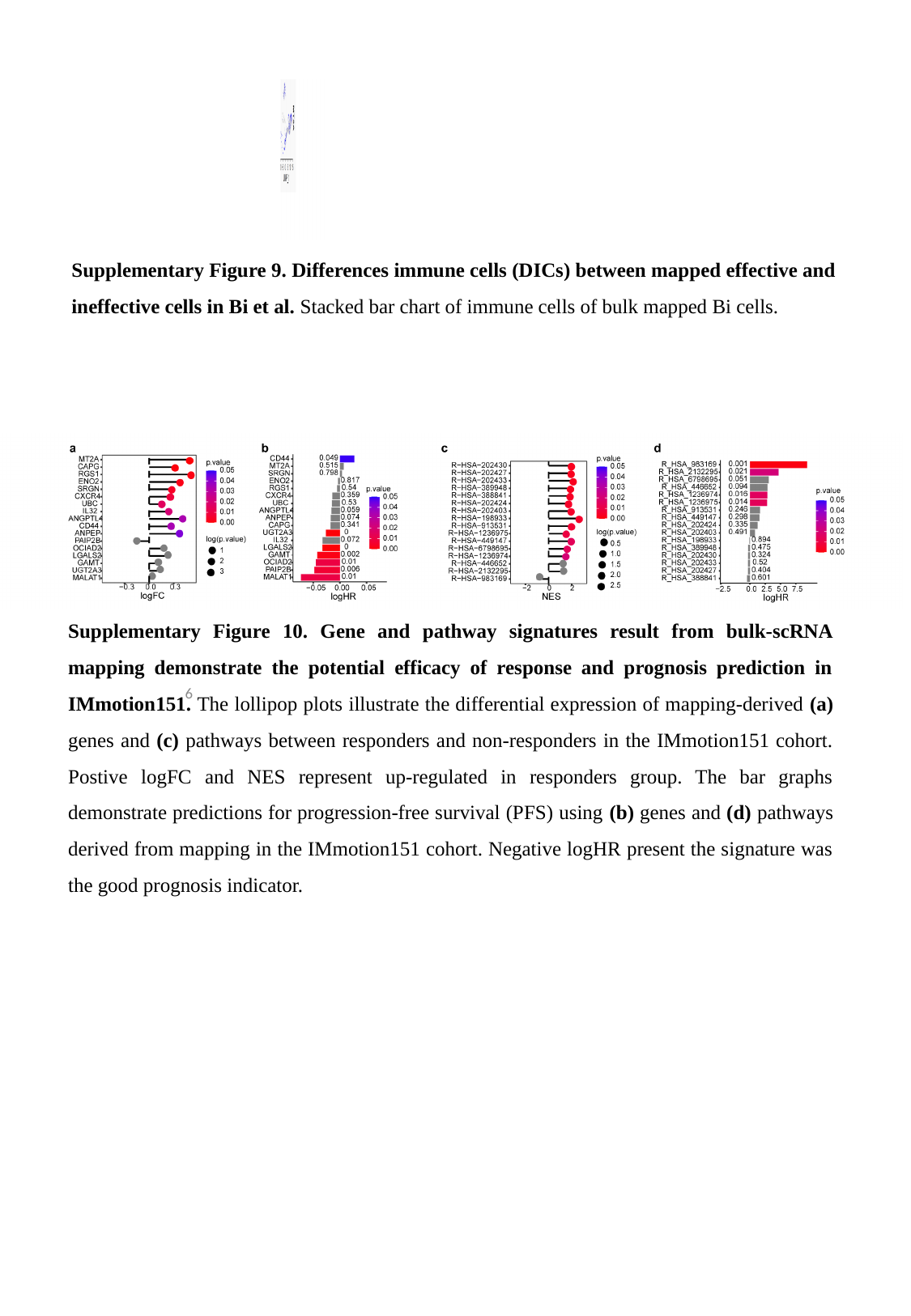

Supplementary Figure 9. Differences immune cells (DICs) between mapped effective and ineffective cells in Bi et al. Stacked bar chart of immune cells of bulk mapped Bi cells.
Supplementary Figure 10. Gene and pathway signatures result from bulk-scRNA mapping demonstrate the potential efficacy of response and prognosis prediction in IMmotion151. The lollipop plots illustrate the differential expression of mapping-derived (a) genes and (c) pathways between responders and non-responders in the IMmotion151 cohort. Postive logFC and NES represent up-regulated in responders group. The bar graphs demonstrate predictions for progression-free survival (PFS) using (b) genes and (d) pathways derived from mapping in the IMmotion151 cohort. Negative logHR present the signature was the good prognosis indicator.
6

### Slide 7
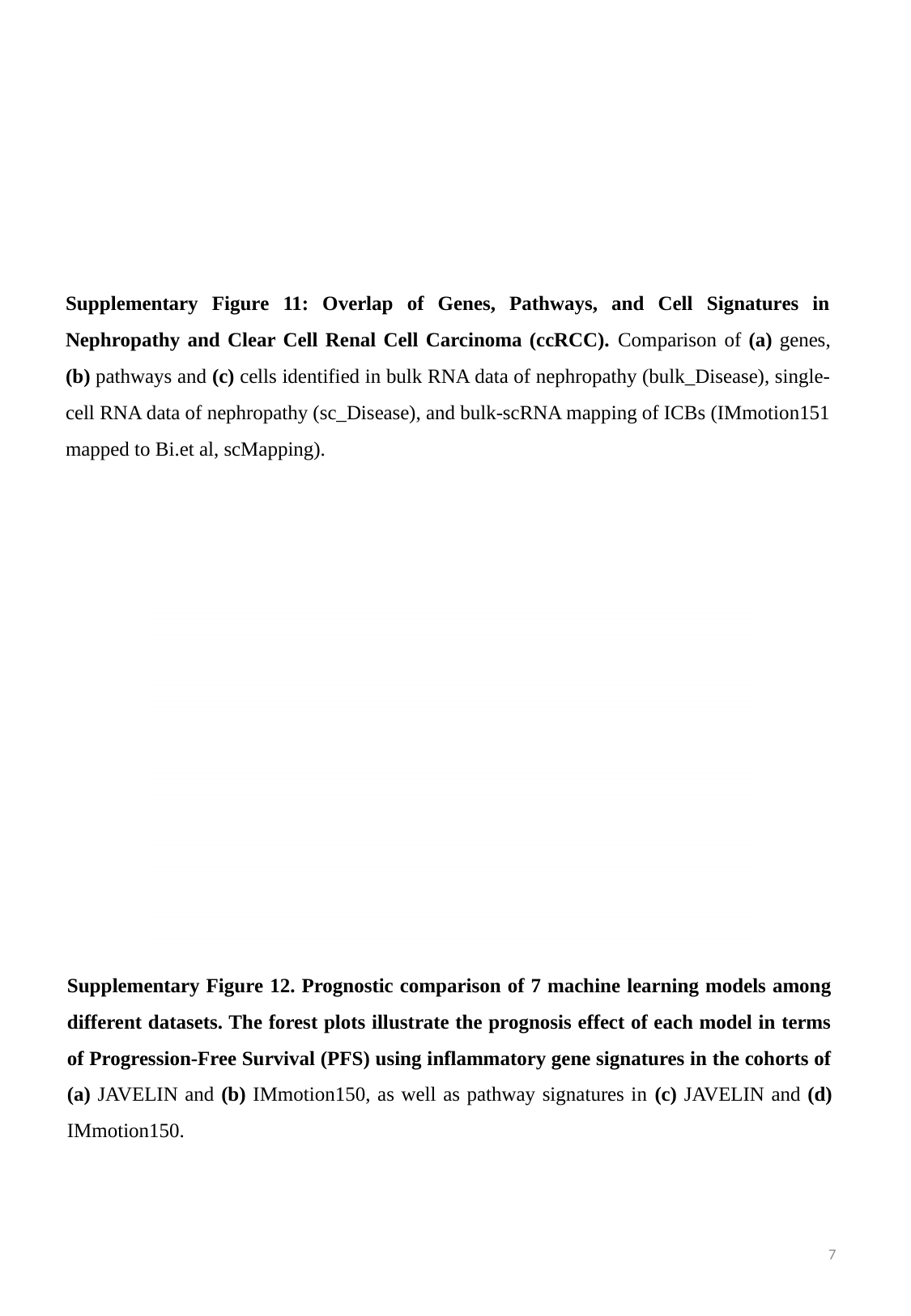

Supplementary Figure 11: Overlap of Genes, Pathways, and Cell Signatures in Nephropathy and Clear Cell Renal Cell Carcinoma (ccRCC). Comparison of (a) genes, (b) pathways and (c) cells identified in bulk RNA data of nephropathy (bulk_Disease), single-cell RNA data of nephropathy (sc_Disease), and bulk-scRNA mapping of ICBs (IMmotion151 mapped to Bi.et al, scMapping).
Supplementary Figure 12. Prognostic comparison of 7 machine learning models among different datasets. The forest plots illustrate the prognosis effect of each model in terms of Progression-Free Survival (PFS) using inflammatory gene signatures in the cohorts of (a) JAVELIN and (b) IMmotion150, as well as pathway signatures in (c) JAVELIN and (d) IMmotion150.
7

### Slide 8
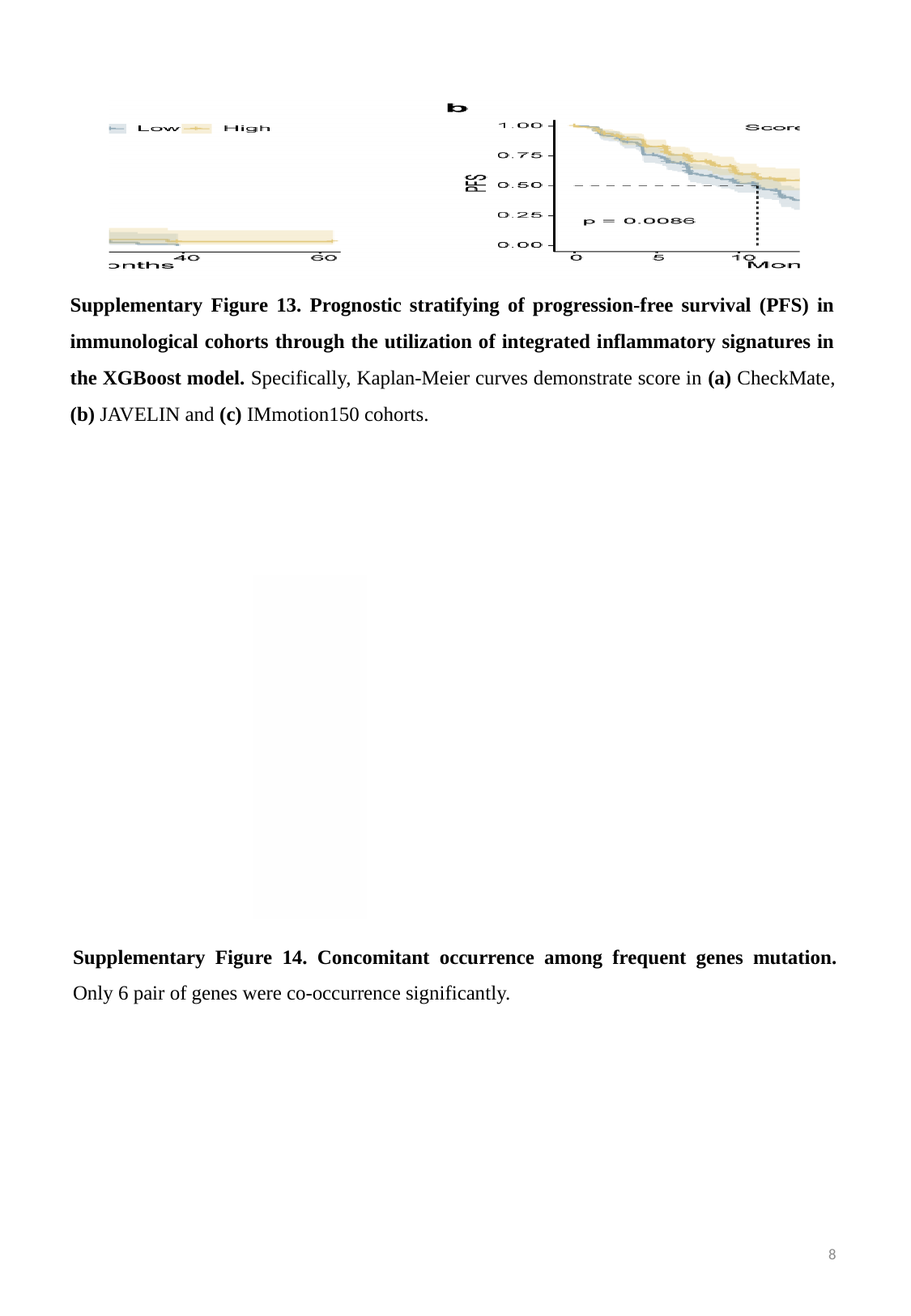

Supplementary Figure 13. Prognostic stratifying of progression-free survival (PFS) in immunological cohorts through the utilization of integrated inflammatory signatures in the XGBoost model. Specifically, Kaplan-Meier curves demonstrate score in (a) CheckMate, (b) JAVELIN and (c) IMmotion150 cohorts.
Supplementary Figure 14. Concomitant occurrence among frequent genes mutation. Only 6 pair of genes were co-occurrence significantly.
8

### Slide 9
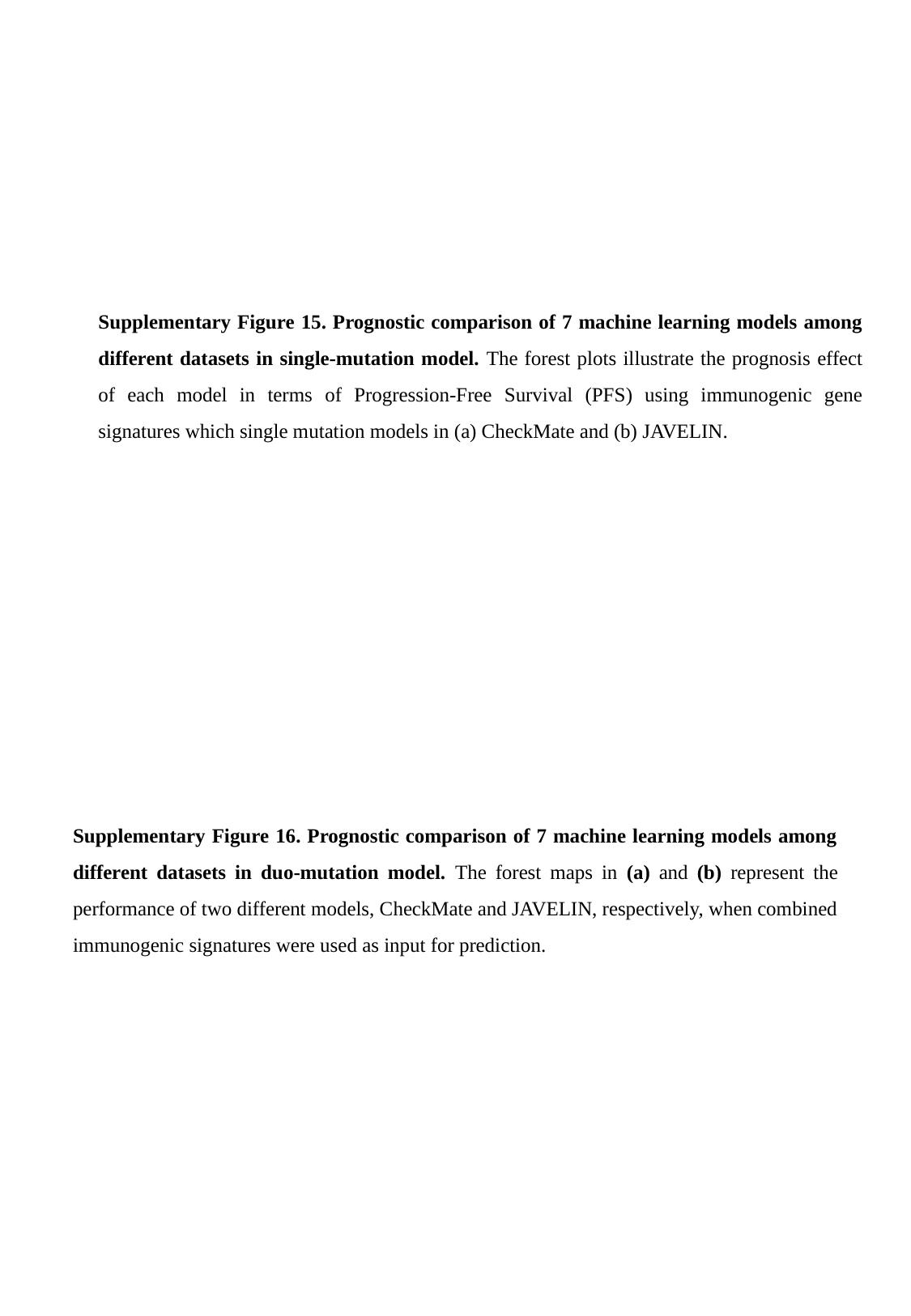

Supplementary Figure 15. Prognostic comparison of 7 machine learning models among different datasets in single-mutation model. The forest plots illustrate the prognosis effect of each model in terms of Progression-Free Survival (PFS) using immunogenic gene signatures which single mutation models in (a) CheckMate and (b) JAVELIN.
Supplementary Figure 16. Prognostic comparison of 7 machine learning models among different datasets in duo-mutation model. The forest maps in (a) and (b) represent the performance of two different models, CheckMate and JAVELIN, respectively, when combined immunogenic signatures were used as input for prediction.

### Slide 10
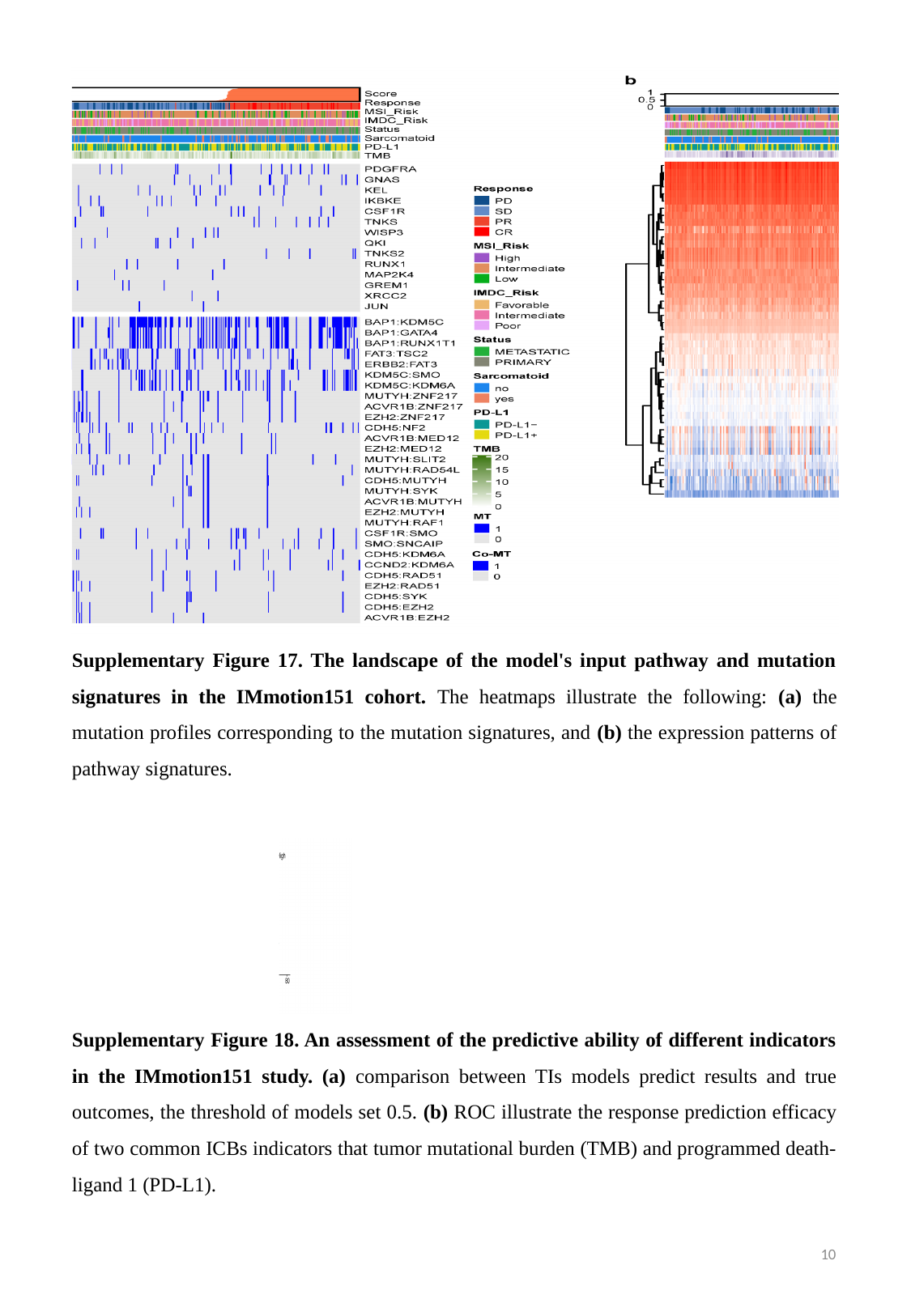

Supplementary Figure 17. The landscape of the model's input pathway and mutation signatures in the IMmotion151 cohort. The heatmaps illustrate the following: (a) the mutation profiles corresponding to the mutation signatures, and (b) the expression patterns of pathway signatures.
Supplementary Figure 18. An assessment of the predictive ability of different indicators in the IMmotion151 study. (a) comparison between TIs models predict results and true outcomes, the threshold of models set 0.5. (b) ROC illustrate the response prediction efficacy of two common ICBs indicators that tumor mutational burden (TMB) and programmed death-ligand 1 (PD-L1).
10

### Slide 11
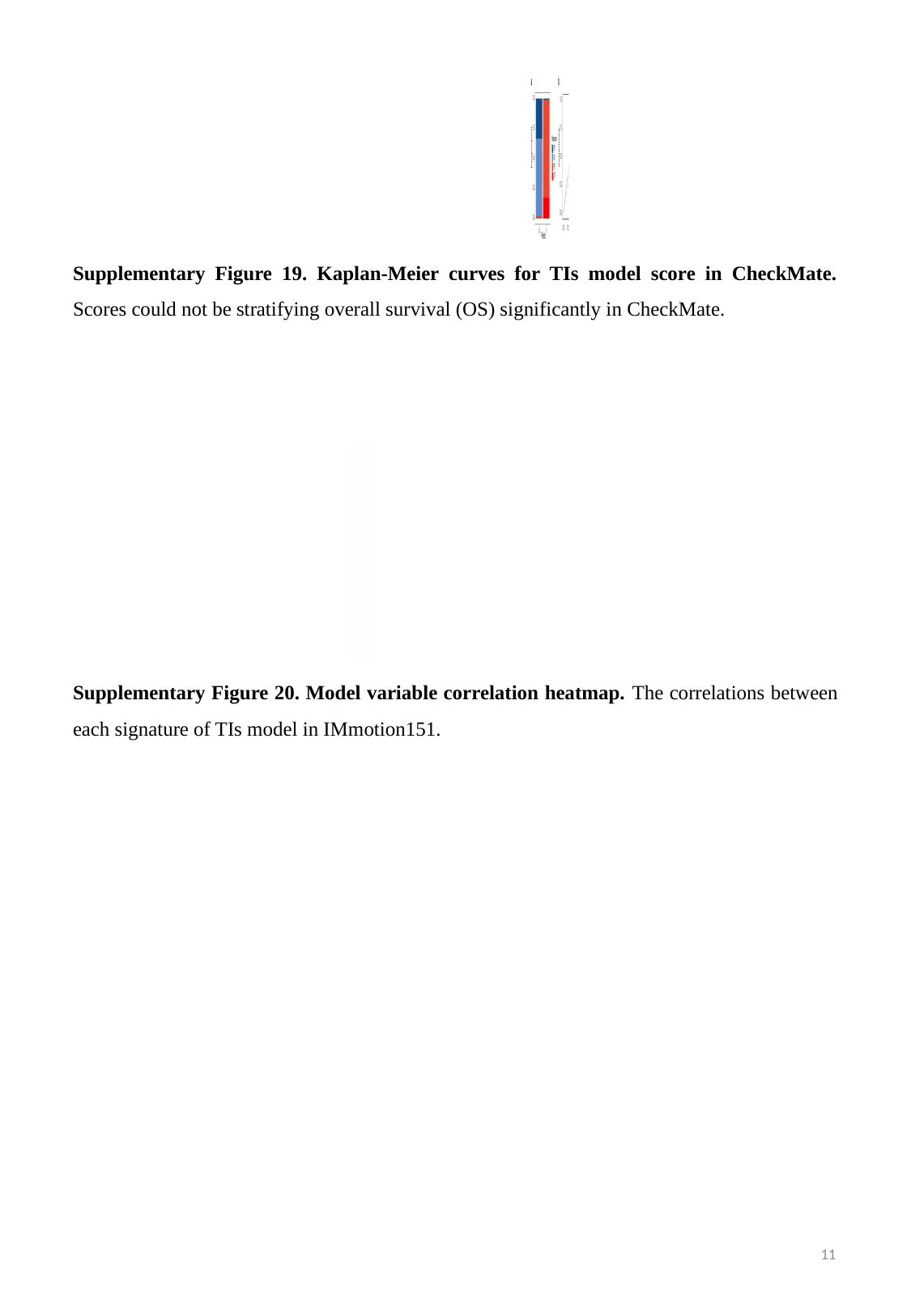

Supplementary Figure 19. Kaplan-Meier curves for TIs model score in CheckMate. Scores could not be stratifying overall survival (OS) significantly in CheckMate.
Supplementary Figure 20. Model variable correlation heatmap. The correlations between each signature of TIs model in IMmotion151.
11

### Slide 12
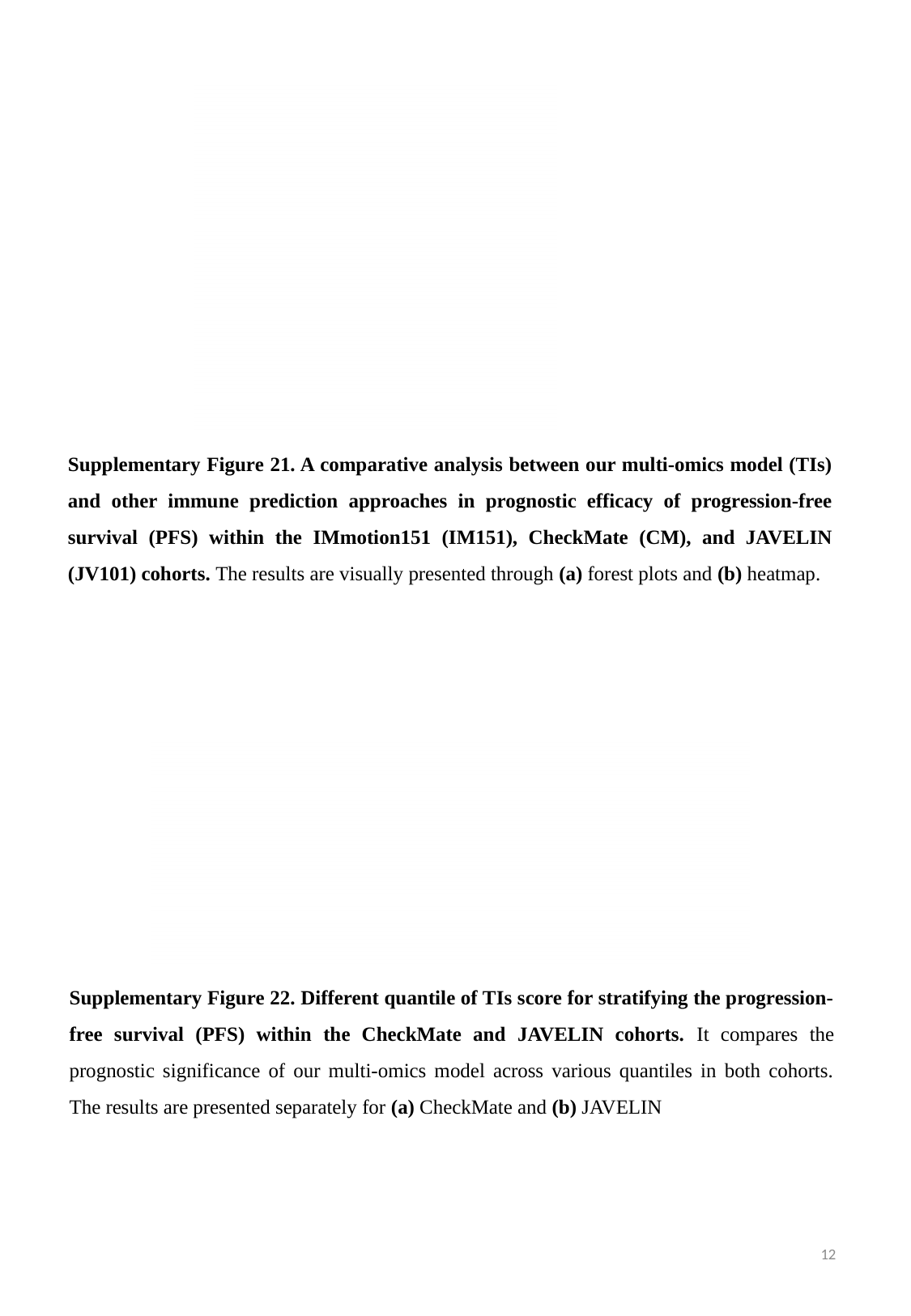

Supplementary Figure 21. A comparative analysis between our multi-omics model (TIs) and other immune prediction approaches in prognostic efficacy of progression-free survival (PFS) within the IMmotion151 (IM151), CheckMate (CM), and JAVELIN (JV101) cohorts. The results are visually presented through (a) forest plots and (b) heatmap.
Supplementary Figure 22. Different quantile of TIs score for stratifying the progression-free survival (PFS) within the CheckMate and JAVELIN cohorts. It compares the prognostic significance of our multi-omics model across various quantiles in both cohorts. The results are presented separately for (a) CheckMate and (b) JAVELIN
12

### Slide 13
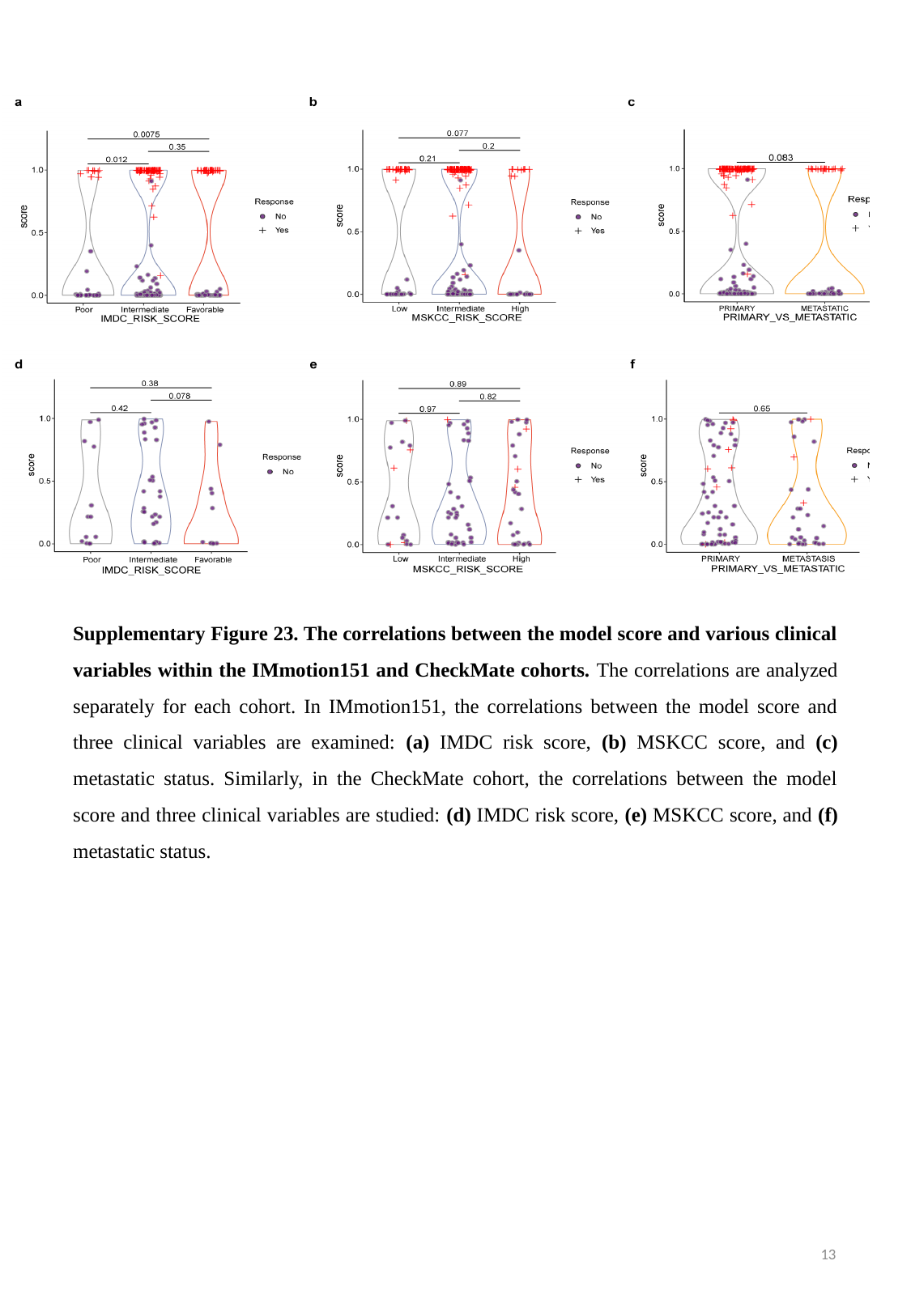

Supplementary Figure 23. The correlations between the model score and various clinical variables within the IMmotion151 and CheckMate cohorts. The correlations are analyzed separately for each cohort. In IMmotion151, the correlations between the model score and three clinical variables are examined: (a) IMDC risk score, (b) MSKCC score, and (c) metastatic status. Similarly, in the CheckMate cohort, the correlations between the model score and three clinical variables are studied: (d) IMDC risk score, (e) MSKCC score, and (f) metastatic status.
13

### Slide 14
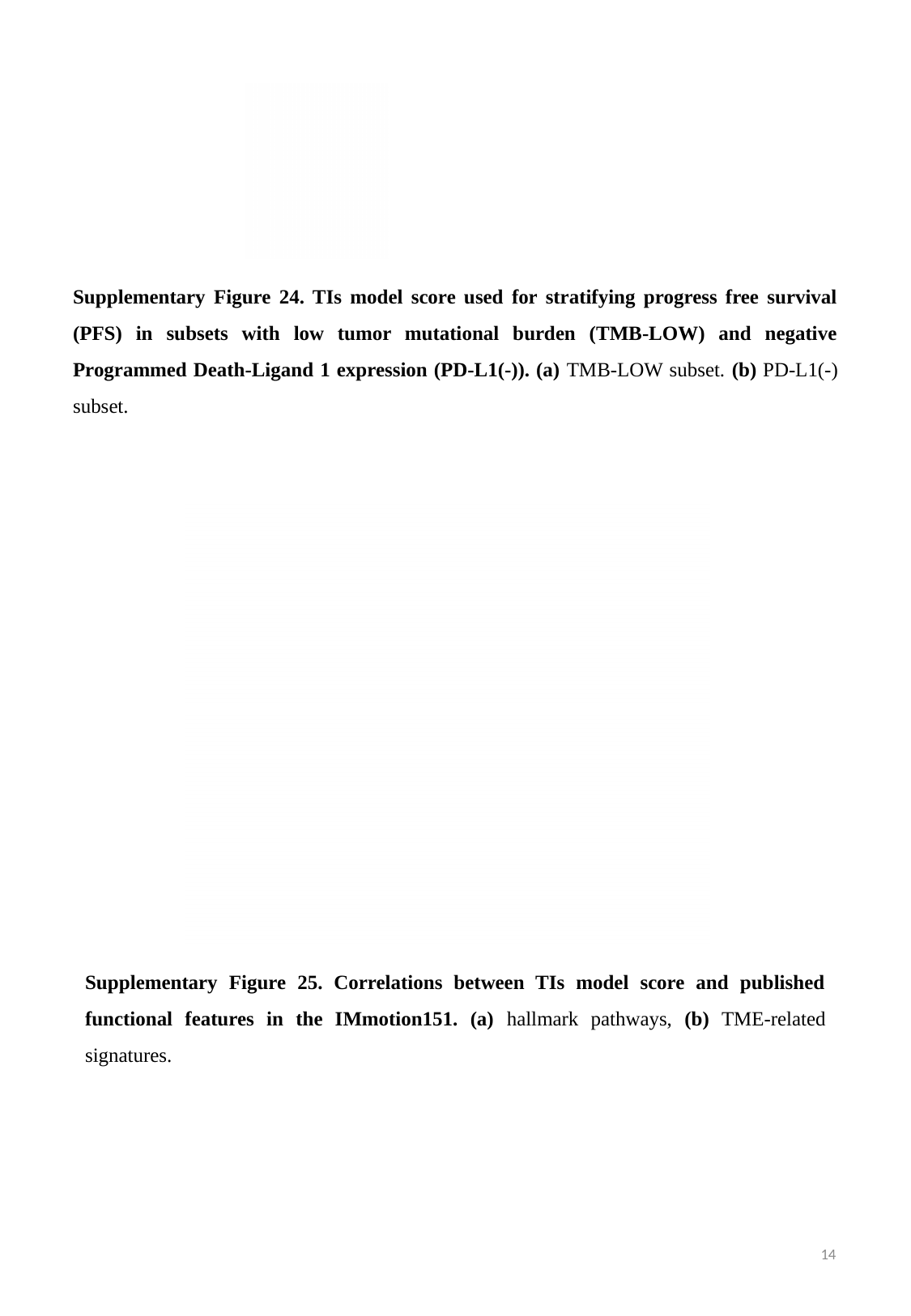

Supplementary Figure 24. TIs model score used for stratifying progress free survival (PFS) in subsets with low tumor mutational burden (TMB-LOW) and negative Programmed Death-Ligand 1 expression (PD-L1(-)). (a) TMB-LOW subset. (b) PD-L1(-) subset.
Supplementary Figure 25. Correlations between TIs model score and published functional features in the IMmotion151. (a) hallmark pathways, (b) TME-related signatures.
14

### Slide 15
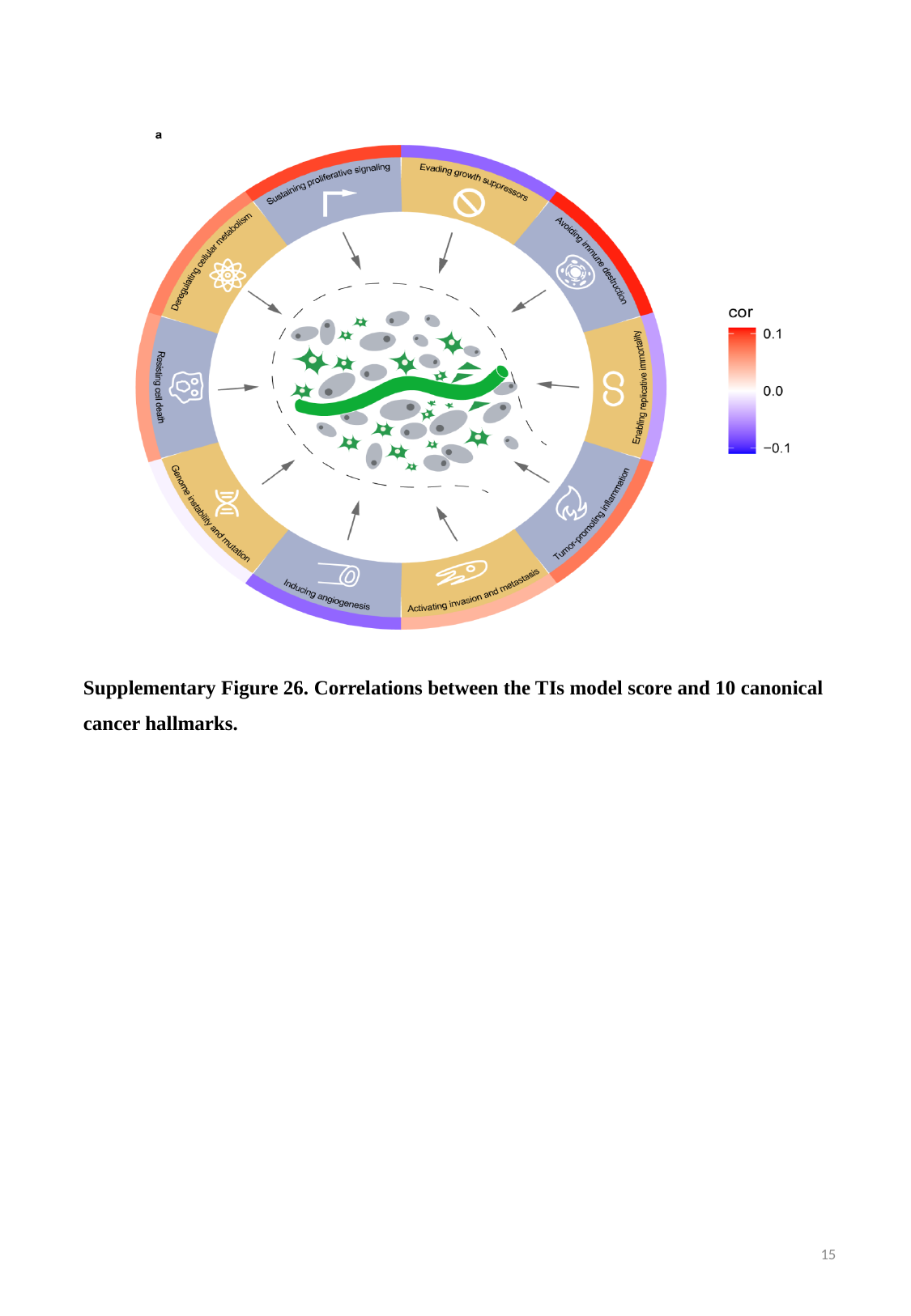

Supplementary Figure 26. Correlations between the TIs model score and 10 canonical cancer hallmarks.
15

### Slide 16
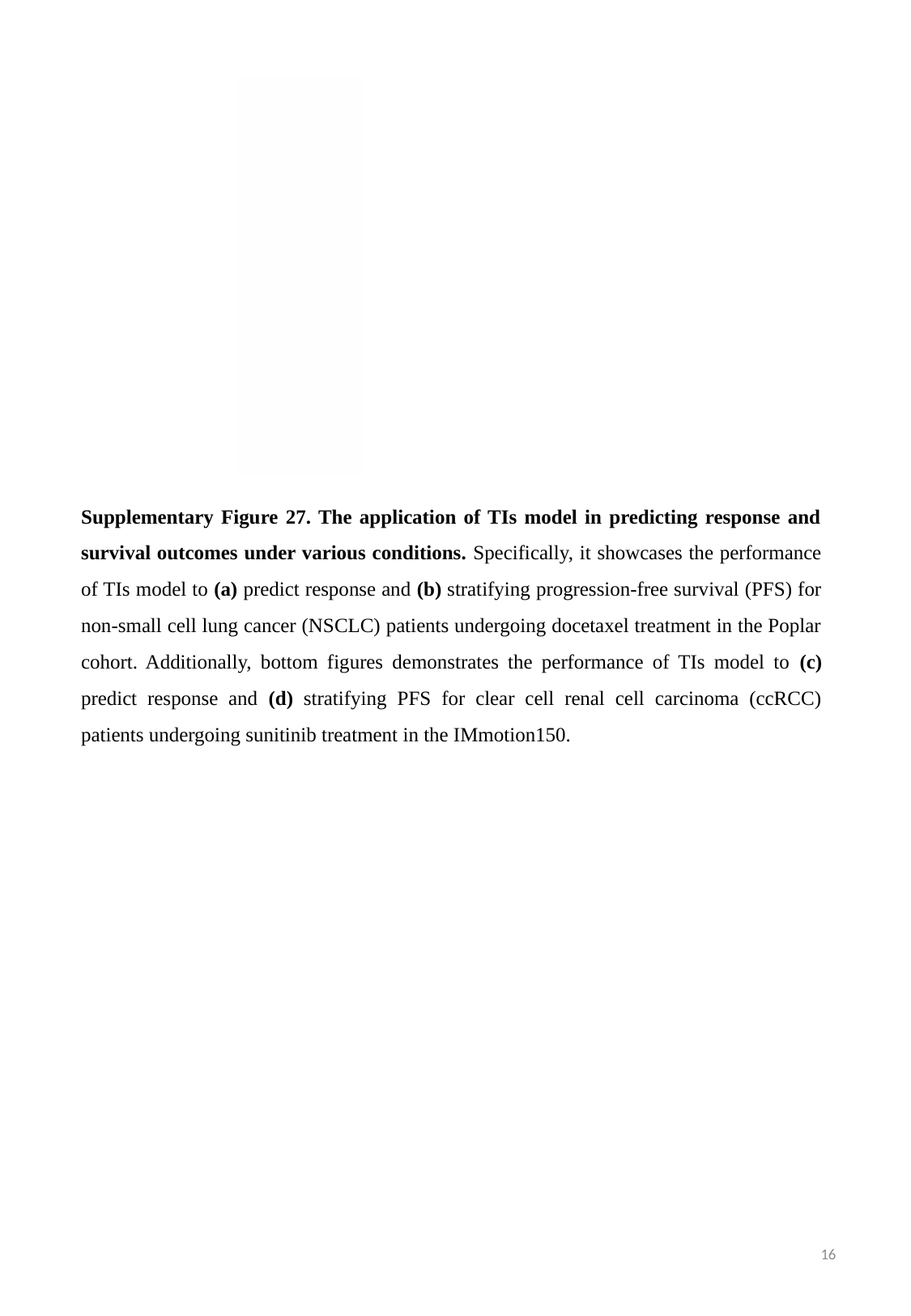

Supplementary Figure 27. The application of TIs model in predicting response and survival outcomes under various conditions. Specifically, it showcases the performance of TIs model to (a) predict response and (b) stratifying progression-free survival (PFS) for non-small cell lung cancer (NSCLC) patients undergoing docetaxel treatment in the Poplar cohort. Additionally, bottom figures demonstrates the performance of TIs model to (c) predict response and (d) stratifying PFS for clear cell renal cell carcinoma (ccRCC) patients undergoing sunitinib treatment in the IMmotion150.
16

### Slide 17
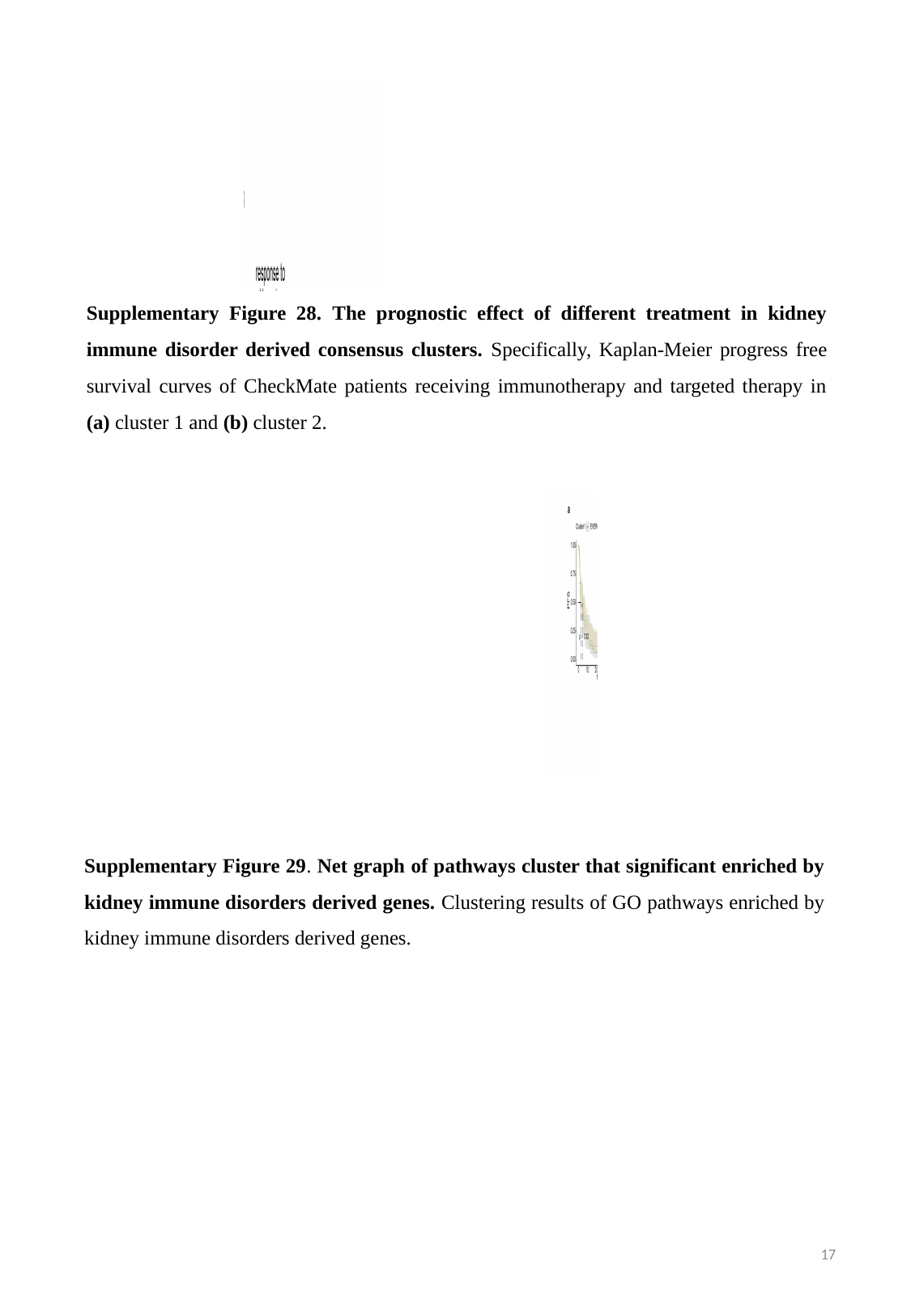

Supplementary Figure 28. The prognostic effect of different treatment in kidney immune disorder derived consensus clusters. Specifically, Kaplan-Meier progress free survival curves of CheckMate patients receiving immunotherapy and targeted therapy in (a) cluster 1 and (b) cluster 2.
Supplementary Figure 29. Net graph of pathways cluster that significant enriched by kidney immune disorders derived genes. Clustering results of GO pathways enriched by kidney immune disorders derived genes.
17

### Slide 18
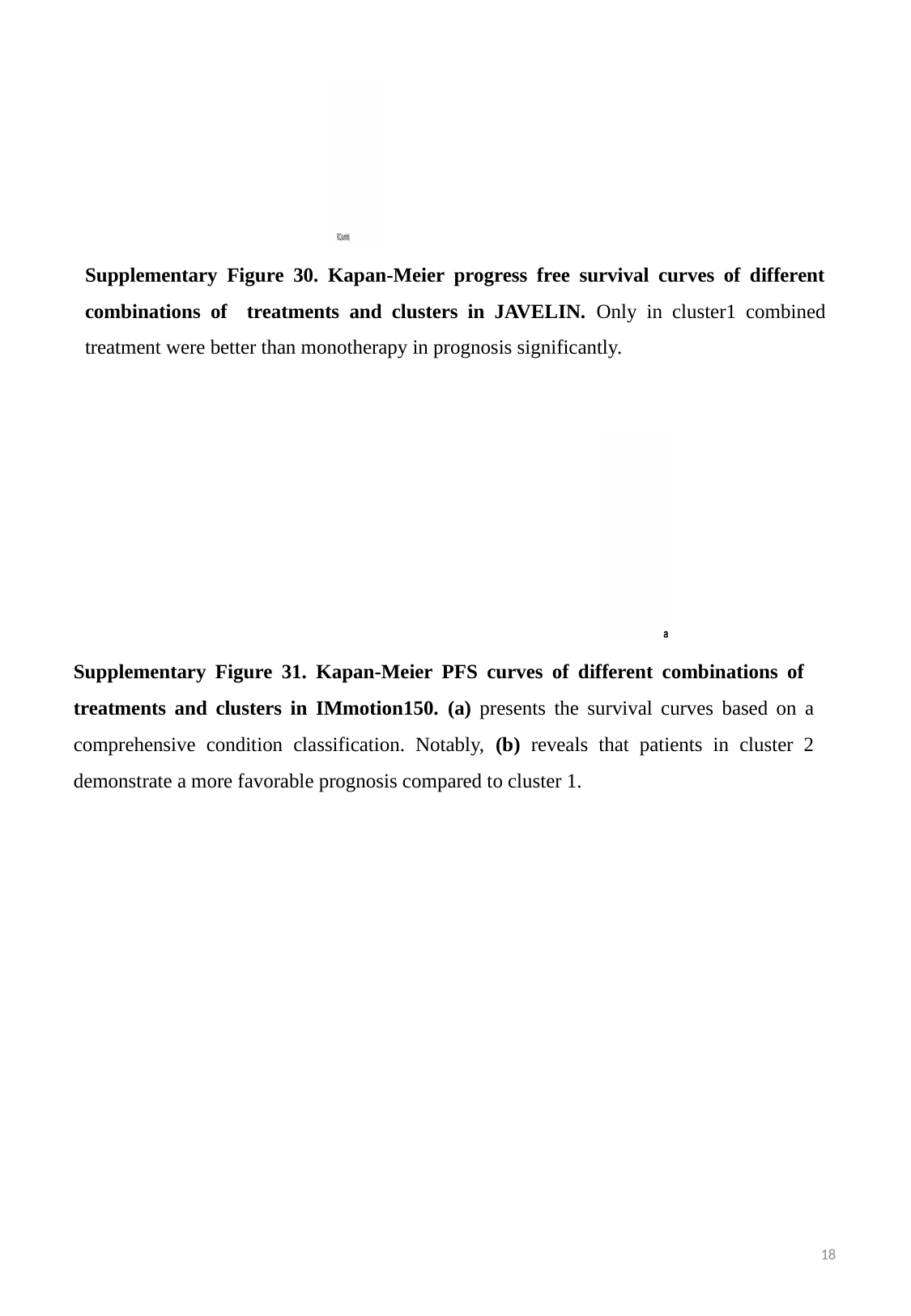

Supplementary Figure 30. Kapan-Meier progress free survival curves of different combinations of treatments and clusters in JAVELIN. Only in cluster1 combined treatment were better than monotherapy in prognosis significantly.
Supplementary Figure 31. Kapan-Meier PFS curves of different combinations of treatments and clusters in IMmotion150. (a) presents the survival curves based on a comprehensive condition classification. Notably, (b) reveals that patients in cluster 2 demonstrate a more favorable prognosis compared to cluster 1.
18

### Slide 19
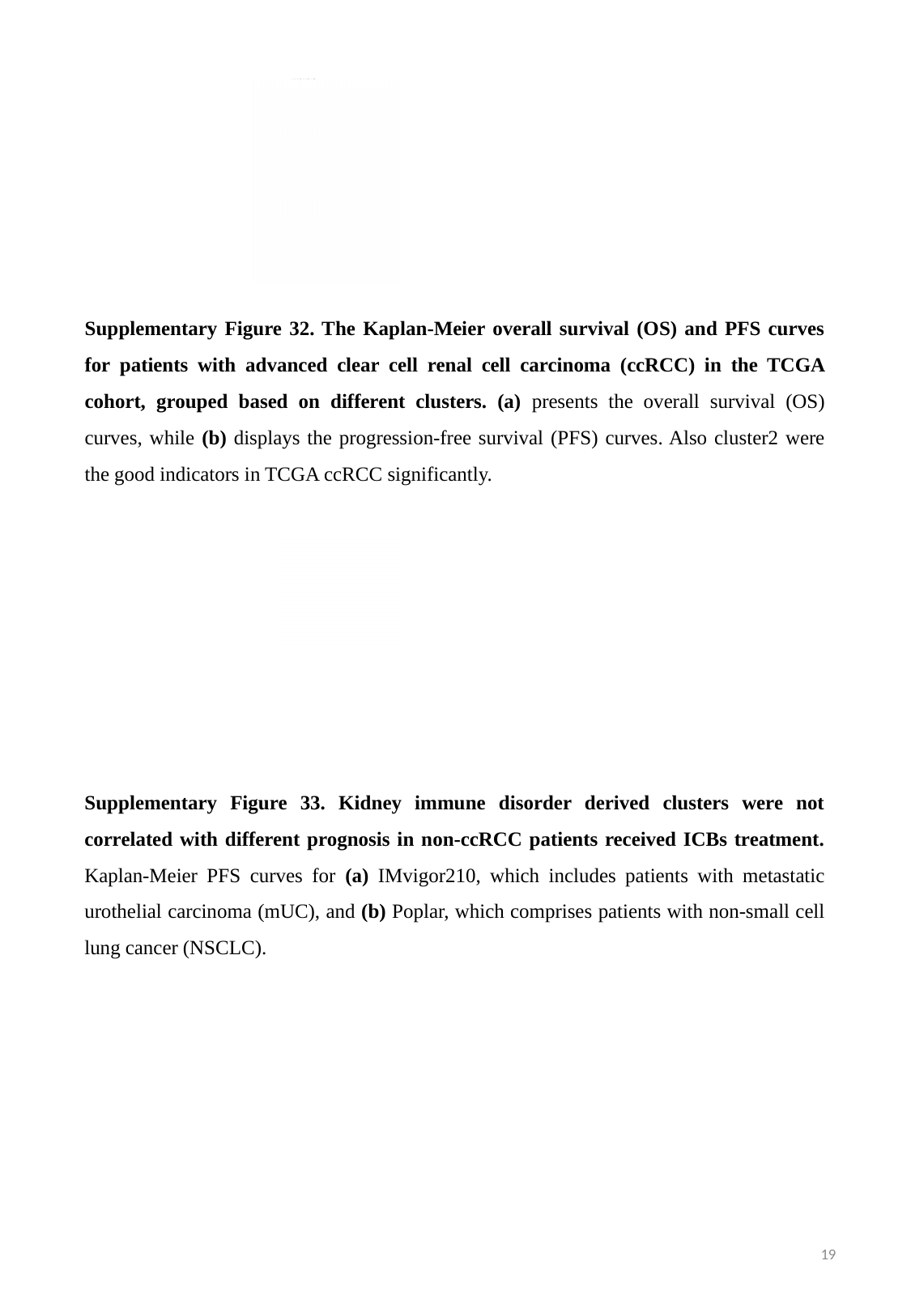

Supplementary Figure 32. The Kaplan-Meier overall survival (OS) and PFS curves for patients with advanced clear cell renal cell carcinoma (ccRCC) in the TCGA cohort, grouped based on different clusters. (a) presents the overall survival (OS) curves, while (b) displays the progression-free survival (PFS) curves. Also cluster2 were the good indicators in TCGA ccRCC significantly.
Supplementary Figure 33. Kidney immune disorder derived clusters were not correlated with different prognosis in non-ccRCC patients received ICBs treatment. Kaplan-Meier PFS curves for (a) IMvigor210, which includes patients with metastatic urothelial carcinoma (mUC), and (b) Poplar, which comprises patients with non-small cell lung cancer (NSCLC).
19
